# Reconstructing A/B compartments as revealed by Hi-C using long-range correlations in epigenetic data

**DOI:** 10.1101/019000

**Authors:** Jean-Philippe Fortin, Kasper D. Hansen

## Abstract

Analysis of Hi-C data has shown that the genome can be divided into two compartments called A/B compartments. These compartments are cell-type specific and are associated with open and closed chromatin. We show that A/B compartments can be reliably estimated using epigenetic data from several different platforms, the Illumina 450k DNA methylation microarray, DNase hypersensitivity sequencing, single-cell ATAC sequencing and single-cell whole-genome bisulfite sequencing. We do this by exploiting the fact that the structure of long range correlations differs between open and closed compartments. This work makes A/B compartments readily available in a wide variety of cell types, including many human cancers.

## Background

Hi-C, a method for quantifying long-range physical interactions in the genome, was introduced by Lieberman-Aiden et al. [2009], and reviewed in Dekker et al. [2013]. A Hi-C assay produces a so-called genome contact matrix which – at a given resolution determined by sequencing depth – measures the degree of interaction between two loci in the genome. In the last 5 years, significant efforts have been made to obtain Hi-C maps at ever increasing resolutions [Dixon et al., 2012, Jin et al., 2013, Naumova et al., 2013, Pope et al., 2014, Rao et al., 2014, Dixon et al., 2015]. Currently, the highest resolution maps are 1kb [Rao et al., 2014]. Existing Hi-C experiments have largely been performed in cell lines or for samples where unlimited input material is available.

In Lieberman-Aiden et al. [2009] it was established that at the megabase scale, the genome is divided into two compartments, called A/B compartments. Interactions between loci are largely constrained to occur between loci belonging to the same compartment. The ‘A’ compartment was found to be associated with open chromatin and the ‘B’ compartment with closed chromatin. Lieberman-Aiden et al. [2009] also showed that these compartments are cell-type specific, but did not comprehensively describe differences between cell types across the genome. In most subsequent work using the Hi-C assay, the A/B compartments have received little attention; the focus has largely been on describing smaller domain structures using higher resolution data. Recently, it was shown that 36% of the genome changes compartment during mammalian development [Dixon et al., 2015] and that these compartment changes are associated with gene expression; they conclude “that the A and B compartments have a contributory but not deterministic role in determining cell-type-specific patterns of gene expression”.

The A/B compartments are estimated by an eigenvector analysis of the genome contact matrix after normalization by the observed-expected method [Lieberman-Aiden et al., 2009]. Specifically, boundary changes between the two compartments occur where the entries of the first eigenvector change sign. The observed-expected method normalizes bands of the genome contact matrix by dividing by their mean. This effectively standardizes interactions between two loci separated by a given distance by the average interaction between all loci separated by the same amount. It is critical that the genome contact matrix is normalized in this way, for the first eigenvector to yield the A/B compartments.

Open and closed chromatin can be defined in different ways using different assays such as DNase hypersensitivity or ChIP sequencing for various histone modifications. While Lieberman-Aiden et al. [2009] established that the ‘A’ compartment is associated with open chromatin profiles from various assays, including DNase hypersensitivity, it was not determined to which degree these different data types measure the same underlying phenomena, including whether the domain boundaries estimated using different assays coincide genomewide.

In this manuscript, we show that we can reliably estimate A/B compartments as defined using Hi-C data by using the Illumina 450k DNA methylation microarray data [Bibikova et al., 2011] as well as DNase hypersensitivity sequencing [Crawford et al., 2006, Boyle et al., 2008]. Both of these data types are widely available on a large number of cell types. In particular the 450k array has been used to profile a large number of primary samples, including many human cancers; more than 20,000 samples are readily available through the Gene Expression Omnibus (GEO) and The Cancer Genome Atlas (TCGA) [TCGA]. We show that our methods can recover cell type differences. This work makes it possible to study A/B compartments comprehensively across many cell types, including primary samples, and to further investigate the relationship between genome compartmentalization and transcriptional activity or other functional readouts. We furthermore show how our methods also works on single-cell epigenetic data, including single-cell assay for transposase-accessible chromatin (ATAC) sequencing [Cusanovich et al., 2015] and single-cell whole-genome bisulfite sequencing (WGBS) [Smallwood et al., 2014].

As an application, we show how the somatic mutation rate in prostate adenocarcinoma is different between compartments and we show how the A/B compartments change between several human cancers; currently The Cancer Genome Atlas does not include assays measuring chromatin accessibility. Furthermore, our work reveals unappreciated aspects of the structure of long-range correlations in DNA methylation and DNase hypersensitivity data. Specifically, we observe that both DNA methylation and DNase signal are highly correlated between distant loci, provided that the two loci are both in the closed compartment.

## Results

### A/B compartments are highly reproducible and are cell-type specific

We obtained publicly available Hi-C data on EBV-transformed lymphoblastoid cell lines (LCLs) and fibroblast cell lines and estimated A/B compartments by an eigenvector analysis of the normalized Hi-C contact matrix (Methods). The contact matrices were preprocessed with ICE [Imakaev et al., 2012] and normalized using the expected-observed method [Lieberman-Aiden et al., 2009]. As in Lieberman-Aiden et al. [2009], we found that the eigenvector divides the genome into two compartments based on the sign of its entries. These two compartments have previously been found to be associated with open and closed chromatin; in the following we will use open to refer to the ‘A’ compartment and closed to refer to the ‘B’ compartment. The sign of the eigenvector is arbitrary; in this manuscript we select the sign so that positive values are associated with the closed compartment (Methods). In Figure 1 we show estimated eigenvectors at 100kb resolution from chromosome 14 across 2 cell types measured in multiple laboratories with widely different sequencing depth, as well as variations in the experimental protocol. We observe a very high degree of correspondence between replicates of the same cell type; on chromosome 14, the correlation between eigenvectors from experiments in the same cell type is greater than 96% (ranges from 96.3% to 98.4%). The agreement, defined as the percentage of genomic bins which are assigned to the same compartment in two different experiments, is greater than 92% (ranges from 92.6% to 96.0%).

**Figure 1.**
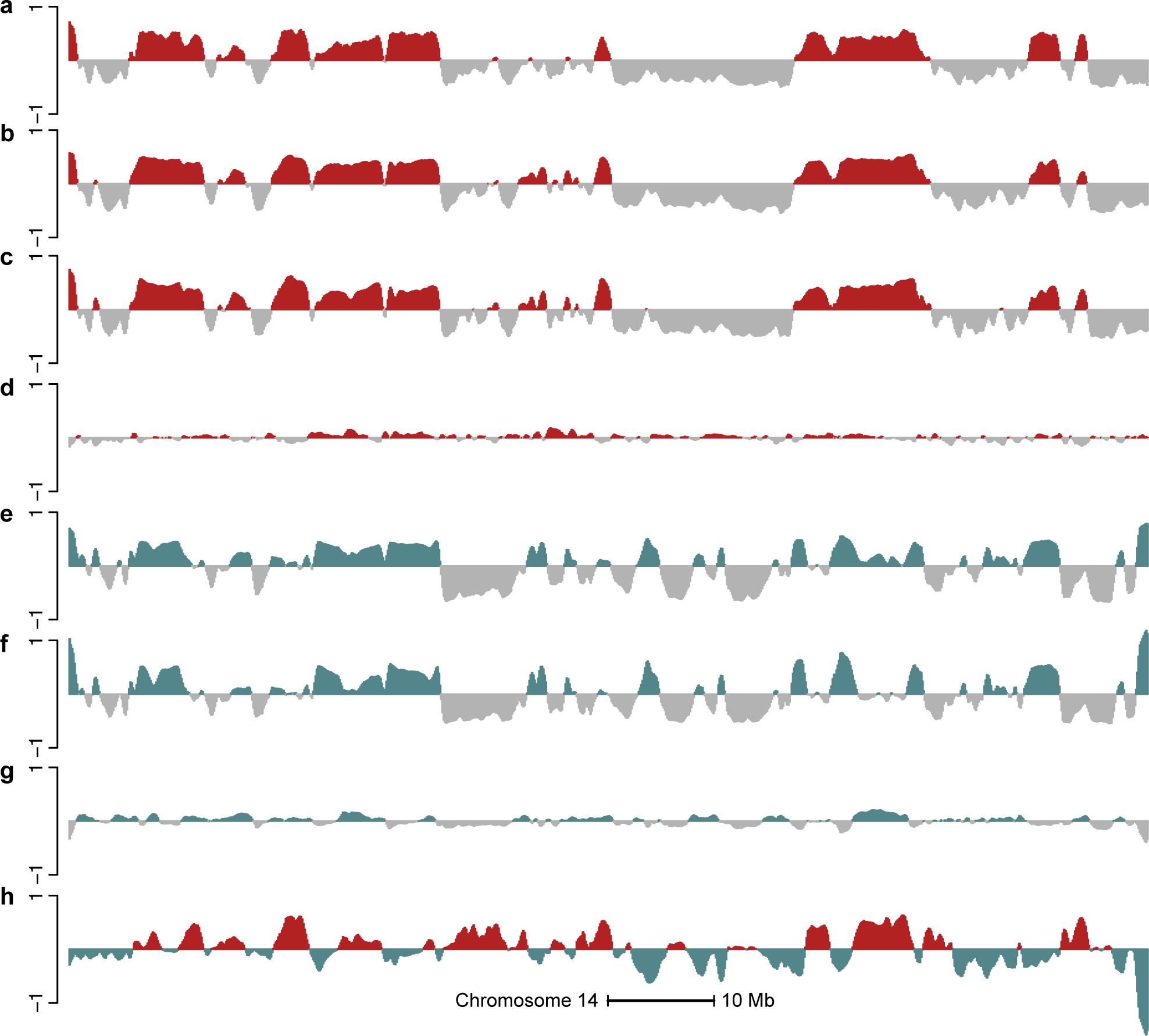
A/B compartments are reproducible and cell type specific. The figure displays data on all of chromosome 14 at 100kb resolution. The first eigenvector for the expected-observed normalized (a) EBV-2009, (b) EBV-2012, (c) EBV-2014 datasets. (d) The difference between (b) and (c). The first eigenvector for the expected-observed normalized (e) IMR-2013, (f) IMR-2014 datasets and (g) their difference. (h) The difference between (c) and (f), which is greater than the technical variation depicted in and (g). This establishes that Hi-C compartments are highly reproducible between experiments in different laboratories and that compartments are cell type specific.

Using high resolution data does not change the estimated A/B compartments as seen in Figure 2.

**Figure 2.**
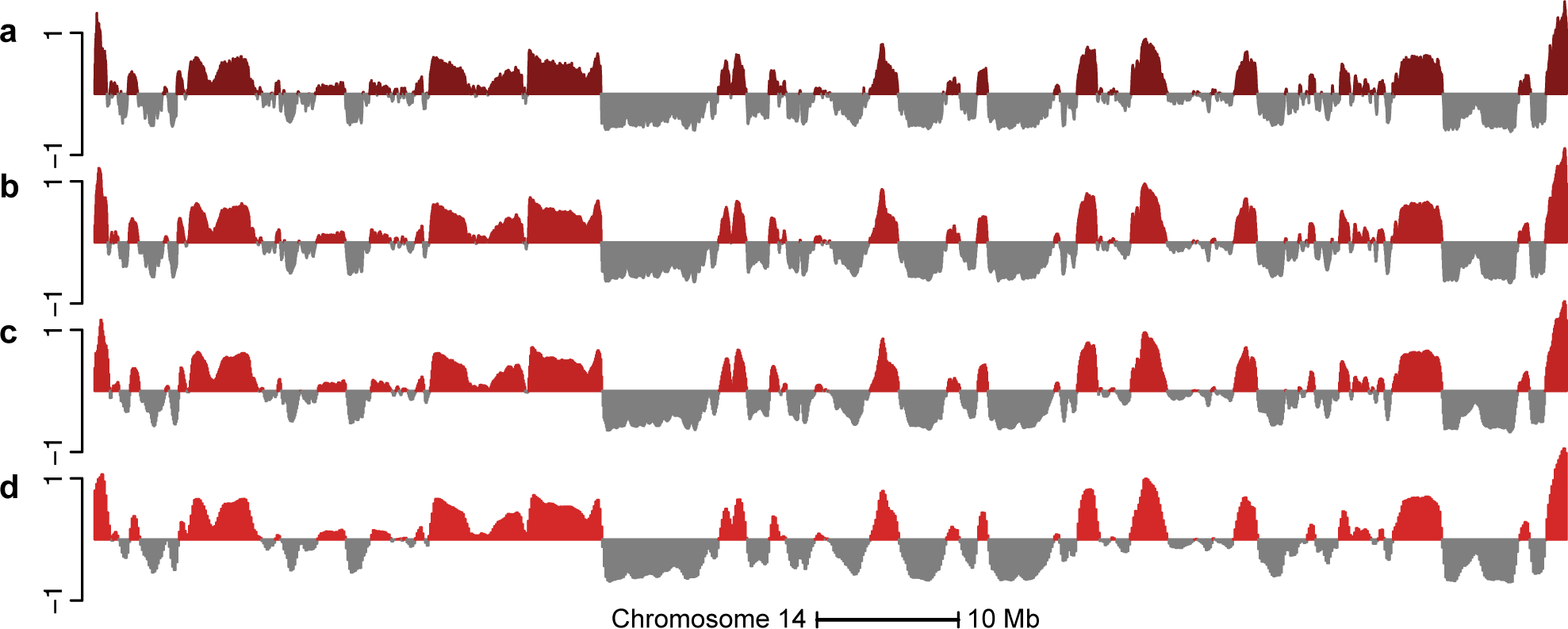
A/B compartments revealed by Hi-C data do not change at resolutions higher than 100kb. The figure displays data on all of chromosome 14 at different resolutions. The four different tracks represent the first eigenvector of the Hi-C IMR90-2014 dataset at resolutions (a) 10kb, (b) 25kb, (c) 50kb and (d) 100kb.

Figure 1 shows the A/B compartments are cell-type specific, with a variation between cell types which exceeds technical variation in the assay; this has been previously noted [Lieberman-Aiden et al., 2009, Dixon et al., 2015]. The correlation between eigenvectors from different cell types is around 60%, in contrast to 96+% between eigenvectors from the same cell type.

In the remainder of the manuscript, we use the most recent data, ie. EBV-2014 and IMR90-2014, to represent eigenvectors and A/B compartments derived from Hi-C data in these cell types.

### Predicting A/B compartments from DNA methylation data

To estimate A/B compartments using other types of epigenetic data we first concentrate on DNA methylation data assayed using the Illumina 450k microarray platform. Data from this platform is widely available across many different primary cell types. To compare with existing Hi-C maps, we obtained data from 288 EBV-transformed lymphoblastoid cell lines from the HapMap project [Heyn et al., 2013].

DNA methylation is often described as related to active and inactive parts of the genome. Most established is high methylation in a genic promoter leading to silencing of the gene [Deaton and Bird, 2011]. As a first attempt to predict A/B compartments from DNA methylation data, we binned the genome and averaged methylation values across samples and CpGs inside each bin. Only CpGs more than 4kb away from CpG islands were used; these are termed open sea CpGs (Methods). We found that high levels of average methylation were associated with the open compartment and not the closed compartment; this might be a consequence of averaging over open sea probes. Figure 3 depicts data from such an analysis for lymphoblastoid cell lines on chromosome 14 at a 100kb resolution and shows some agreement between estimated compartments from Hi-C and this analysis, with a correlation of 56.3% and a compartment agreement between datasets of 71.7%. In this analysis, we implicitly assume that there is no variation in compartments between different individuals in the same cell type.

**Figure 3.**
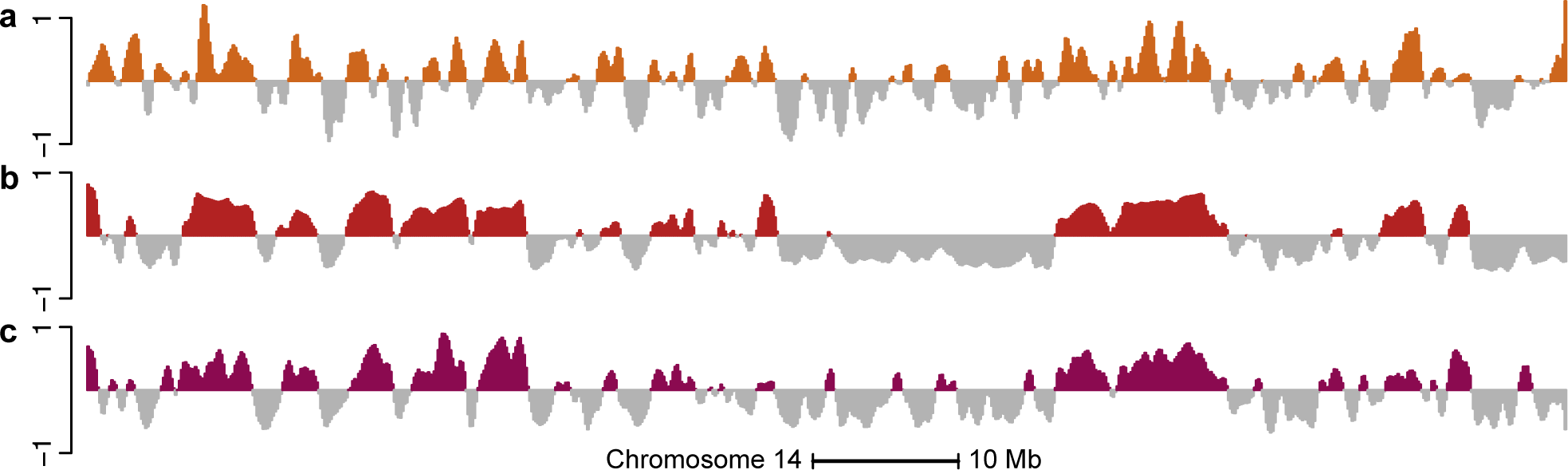
The methylation correlation signal is a better predictor of A/B compartments than the average methylation signal. The figure displays data on all of chromosome 14 at 100kb resolution. (a) The smoothed, average methylation signal on the beta-value scale for the 450k-EBV dataset. The signal has been centered by the mean and the sign has been reversed so that values close to one correspond to low methylation values. (b) The first eigenvector of the EBV-2014 Hi-C dataset. (c) The smoothed first eigenvector of the binned correlation matrix of the 450k-EBV dataset. We see that (c) correlates better with (b) than (a).

Surprisingly, we found that we could improve considerably on this analysis by doing an eigenvector analysis of a suitably processed between-CpG correlation matrix (Figure 3). This matrix represents correlations between any two CpGs measured on the 450k array, with the correlation being based on biological replicates of the same cell type. The correlation eigenvector shows strong agreement with the Hi-C eigenvector; certainly higher than with the average methylation vector (Figure 3). Quantifying this agreement, we find that the correlation between the two vectors is 84.8% and the compartment agreement is 83.8% on chromosome 14. Genomewide, the correlation is 70.9% and the agreement is 79% (Table 1); we tend to perform worse on smaller chromosomes. Again, this analysis implicitly assumes lack of variation in compartments between biological replicates.

**Table 1.** Correlation and agreement between Hi-C and 450k-based eigenvector estimates of genome compartments. Thresholding refers to excluding genomic bins where the entries of the relevant eigenvector has an absolute value less than 0.01

|  | Chr 14 |  | Genome |  |
| --- | --- | --- | --- | --- |
|  | EBV | Fibro | EBV | Fibro |
| No threshold |  |  |  |  |
| Correlation | 84.8% | 75.1% | 70.0% | 73.6% |
| Agreement | 83.7% | 79.1% | 79.0% | 79.5% |
| Bins retained | 100% | 100% | 100% | 100% |
| Threshold, Methylation |  |  |  |  |
| Correlation | 87.4% | 78.4% | 74.2% | 77.3% |
| Agreement | 88.8% | 83.4% | 87.9% | 87.9% |
| Bins retained | 86% | 85% | 78% | 79% |
| Threshold, Methylation and Hi-C |  |  |  |  |
| Correlation | 89.1% | 84.2% | 77.4% | 81.1% |
| Agreement | 93.0% | 90.5% | 92.6% | 92.8% |
| Bins retained | 76% | 67% | 66% | 64% |

Closely examining differences between the 450k-based predictions and the Hi-C-based estimates, we find that almost all disagreements between the two methods occur when entries in one of the two eigenvectors is close to zero; in other words, where there is uncertainty about the compartment in either of the two analyses. Excluding bins where the 450k-based prediction is close to zero, that is bins that have an absolute eigenvector value less than 0.01, we get an agreement of 88.8% (14.2% of the bins excluded). Excluding bins where either the 450k-based prediction is close to zero or the Hi-C eigenvector is close to zero, we get an agreement of 93% (24.8% of the bins excluded).

Our processing of the correlation matrix is as follows (details are in Methods and rationale is explained below): in our correlation matrix, we only include so-called ‘open sea’ CpGs; these CpGs are more than 4kb away from CpG islands. Next, we bin each chromosome into 100kb bins and compute which open sea CpGs are inside each bin; this will vary between bins due to the design of the 450k microarray. To get a single number representing the correlation between two bins, we take the median of the correlations of the individual CpGs located in each bin. We obtain the first eigenvector of this binned correlation matrix and gently smooth the signal by using two iterations of a moving average with a window size of 3 bins.

The sign of the eigenvector is chosen so that the sign of the correlation between the eigen-vector and column sums of the correlation matrix is positive; this ensures that positive values of the eigenvector is associated with the closed compartment (see Methods).

### Long-range correlations in DNA methylation data predicts A/B compartment changes between cell types

To examine how well the predictions based on long-range correlations in 450k data captures differences between cell types, we obtained publicly available 450k data from 62 fibroblast samples [Wagner et al., 2014], and compared them to Hi-C data from the IMR90 cell lines. Note that the fibroblast cell lines assayed on the 450k platform are from primary skin in contrast to the IMR90 cell line, a fetal lung fibroblast. Figure 4 and Table 1 shows our ability to recover the A/B compartments in fibroblasts; it is similar to our performance for EBV-transformed lymphocytes.

**Figure 4.**
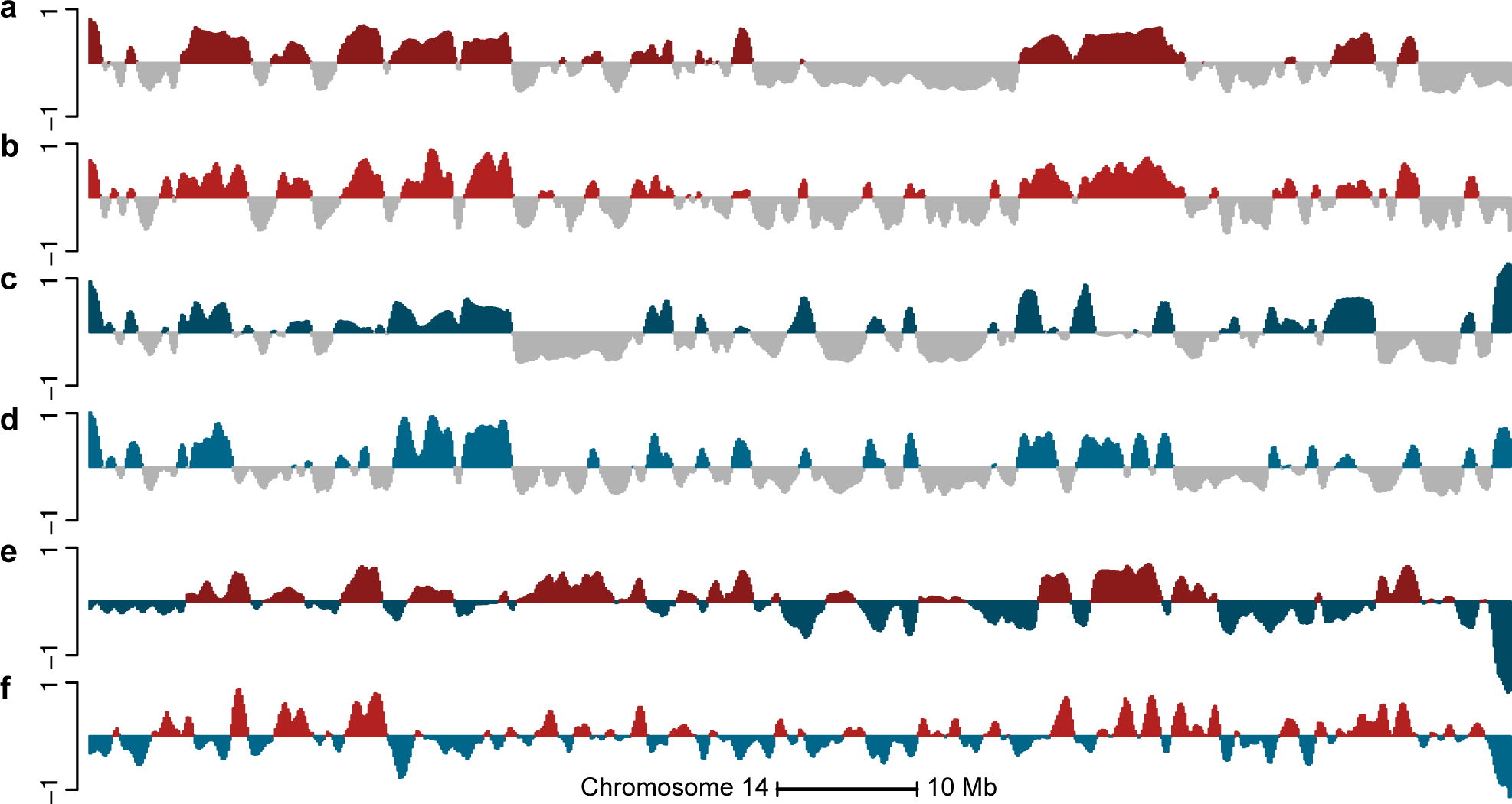
Cell-type specific A/B compartments using Hi-C data are predicted using DNA methylation data. The figure displays data on all of chromosome 14 at 100kb resolution. (a) The first eigenvector of the EBV-2014 Hi-C dataset. (b) The smoothed first eigenvector of the binned correlation matrix of the 450k-EBV dataset. (c) The first eigenvector of the IMR90-2014 Hi-C dataset. (d) The smoothed first eigenvector of the binned correlation matrix of the 450k-Fibroblast dataset. (e) The difference between (a) and (c). (f) the difference between (b) and (d). The high correlation between (e) and (f) supports that the correlation eigenvectors of the 450k data can be used to find differences between compartments in the two cell types.

To firmly establish that the high correlation between our predicted compartments using DNA methylation and Hi-C data is not due to chance, we compared the predicted compartments in EBV transformed lymphocytes and fibroblasts to Hi-C data from different cell types, including the K562 cell line which serves as a somewhat independent negative control. In Figure 5, we show the correlation and agreement between the two sets of predicted compartments and Hi-C data from the three cell types. There is always a decent agreement between predicted compartments of any two cell types, but the agreement is consistently higher when the prediction is from data from the same cell type as the Hi-C data.

**Figure 5.**
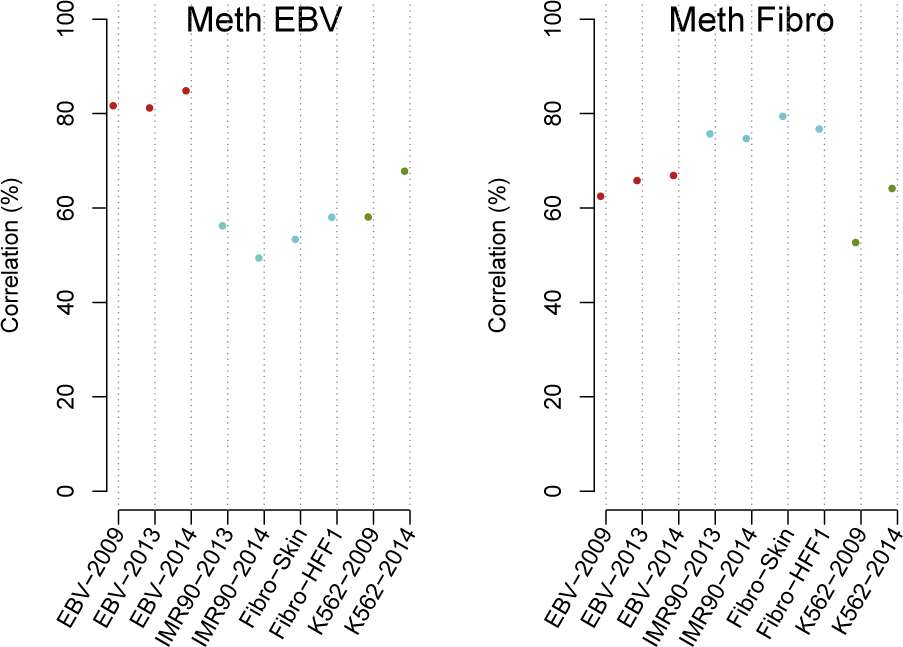
Compartment predictions based on 450k data are cell type specific. Correlations between the eigenvectors from 9 different Hi-C datasets across 3 different cell types with both the 450k-EBV and 450k-Fibroblast datasets. We observe higher correlation between Hi-C data and DNA methylation data when the comparison is being made for the same cell type.

How to best quantify differences in A/B compartments is still an open question. Lieberman-Aiden et al. [2009] used 0 as a threshold to differentiate the two compartments. Considering the difference of two eigenvectors derived in different cell types, it is not clear that functional differences exist exactly when the two eigenvectors have opposite signs; instead, functional differences might be associated with changes in the magnitude of the eigenvectors reflecting a genomic region being relatively more open or closed. We note that the genomic region highlighted as cell-type specific, and validated by FISH, in Lieberman-Aiden et al. [2009], is far away from zero in one condition and has small values fluctuating around zero in the other condition.

Following this discussion, we focus on estimating the direction of change in eigenvectors between different cell types. Figure 4 shows estimated differences between Hi-C and 450k eigenvectors for two cell types. Large differences between the two vectors are replicated well between the two data types, but there is disagreement when the eigenvectors are close to zero. This is to be expected; there is technical variation in such a difference even between Hi-C experiments (Figure 1). Using the data displayed in Figure 1 we find that the technical variation in the Hi-C data is such that 98% of genomic bins have an absolute value less than 0.02. Using this cutoff for technical variation, we find that the correlation between the two difference vectors displayed in Figure 4 is 85% when restricted to the 24% of genomic bins where both vectors have an absolute value greater than 0.02. The sign of the differential vectors are also in high agreement; they agree in 90% of the genomic bins exceeding the cutoff for technical variation. In contrast, the correlation is 61% when the entire chromosome is included, reflecting that the technical noise is less correlated than the signal.

Large domains of intermediate methylation have been previously described [Lister et al., 2009], as well as long blocks of hypomethylation associated with colon cancer and EBV transformation [Berman et al., 2012, Hansen et al., 2011, 2014]. We obtained previously characterized [Lister et al., 2009] partially methylated domains (PMDs) in IMR90 and found a significant overlap with closed compartments from the IMR90-2014 Hi-C dataset (odds ratio: 13.6) as well as closed compartments from the 450k-Fibroblast dataset (odds ratio: 16.4). Likewise, we obtained previously characterized blocks of hypomethylation associated with EBV transformation [Hansen et al., 2014] and found a significant overlap with closed compartments from the EBV-2014 Hi-C dataset (odds ratio: 11.9) and 450k-EBV dataset (odds ratio: 9.4). This confirms previously described overlap between Hi-C compartments and these types of methylation domains by Berman et al. [2012].

### The structure of long-range correlations in DNA methylation data

To understand why we are able to predict open and closed compartments using the 450k array, we studied the structure of long-range correlations in DNA methylation data. First, we note that entries in our binned correlation matrix (within a chromosome) do not decay with distance between bins (Figure 6a). This is in contrast to a Hi-C contact matrix, which has repeatedly been shown to decay with distance as expected (Figure 6b). However, for the first eigenvector to define open and closed compartments, the Hi-C contact matrix needs to be normalized using the expected-observed method [Lieberman-Aiden et al., 2009]. This normalization has the consequence that values in the matrix no longer decay with distance (Figure 6c).

**Figure 6.**
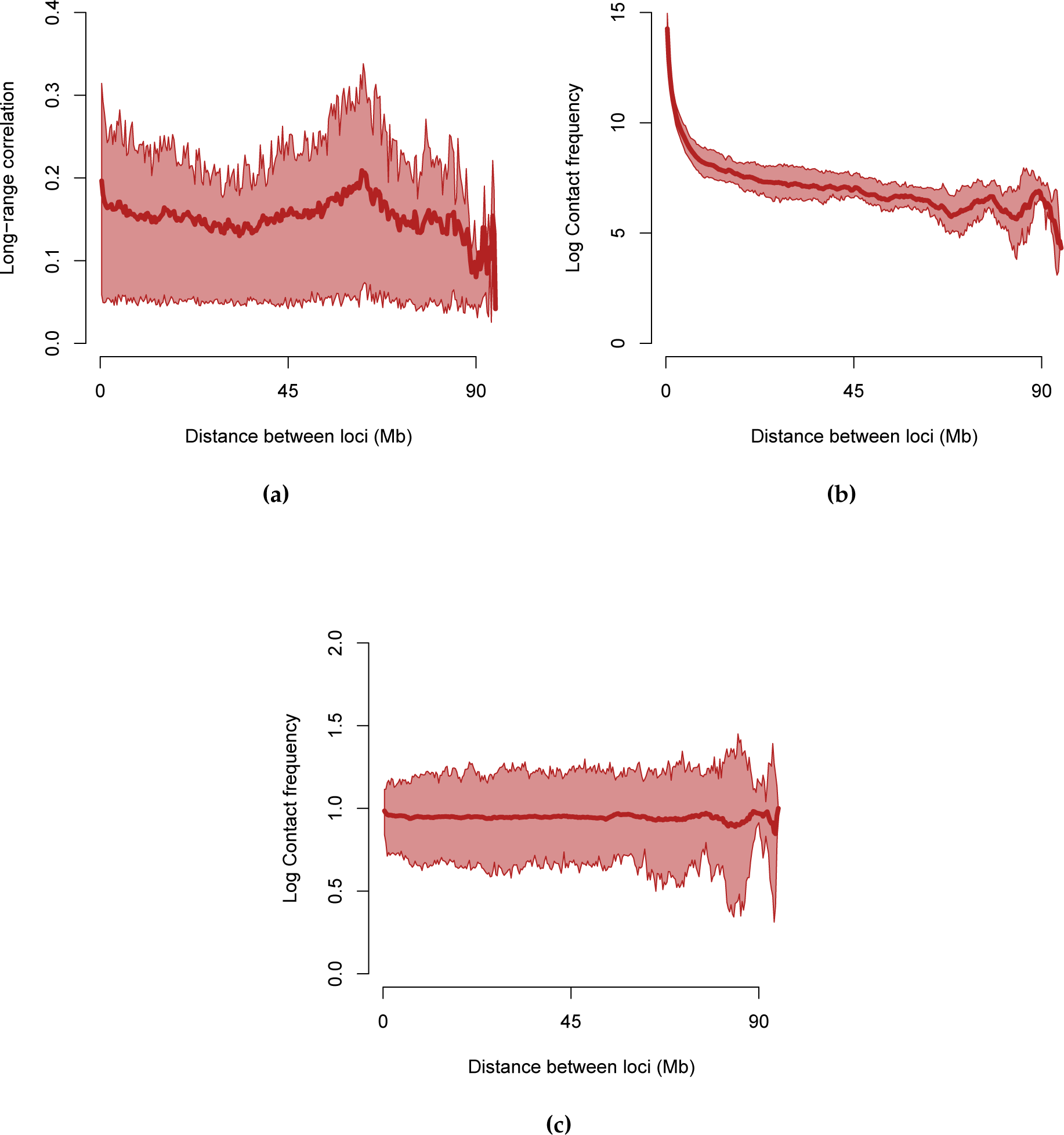
Correlation as function of distance between loci. In each figure we display the median and 25%,75%-quantiles of the entries in a given matrix as a function of genomic distance. (a) Entries in the binned correlation matrix of the 450k-EBV dataset. Entries in the ICE normalized log contact matrix for the EBV-2014 Hi-C experiment. As (b) but the matrix has been normalized using the expected-observed method.

In Figure 7 we show density plots of binned correlations on chromosome 14, stratified in two ways. The first stratification separates correlations between bins which are both in the open compartment, both in the closed and finally cross-compartment correlations. This stratification shows that we have a large amount of intermediate correlation values (0.2-0.5), but only between bins which are both in the closed compartment. The second stratification separates open sea probes and CpG resort probes (probes within 4kb of a CpG island, see Methods). This stratification shows that we only have intermediate correlation values for open sea probes; CpG resort probes are generally uncorrelated. In conclusion, we have the following structure of the binned correlation matrix: most of the matrix contains correlation values around zero (slightly positive), except between two bins both in the closed compartment, which have an intermediate correlation value of 0.2-0.5. This shows why an eigen analysis of the binned correlation matrix recovers the open and closed compartments, see Figure 8 for an illustration.

**Figure 7.**
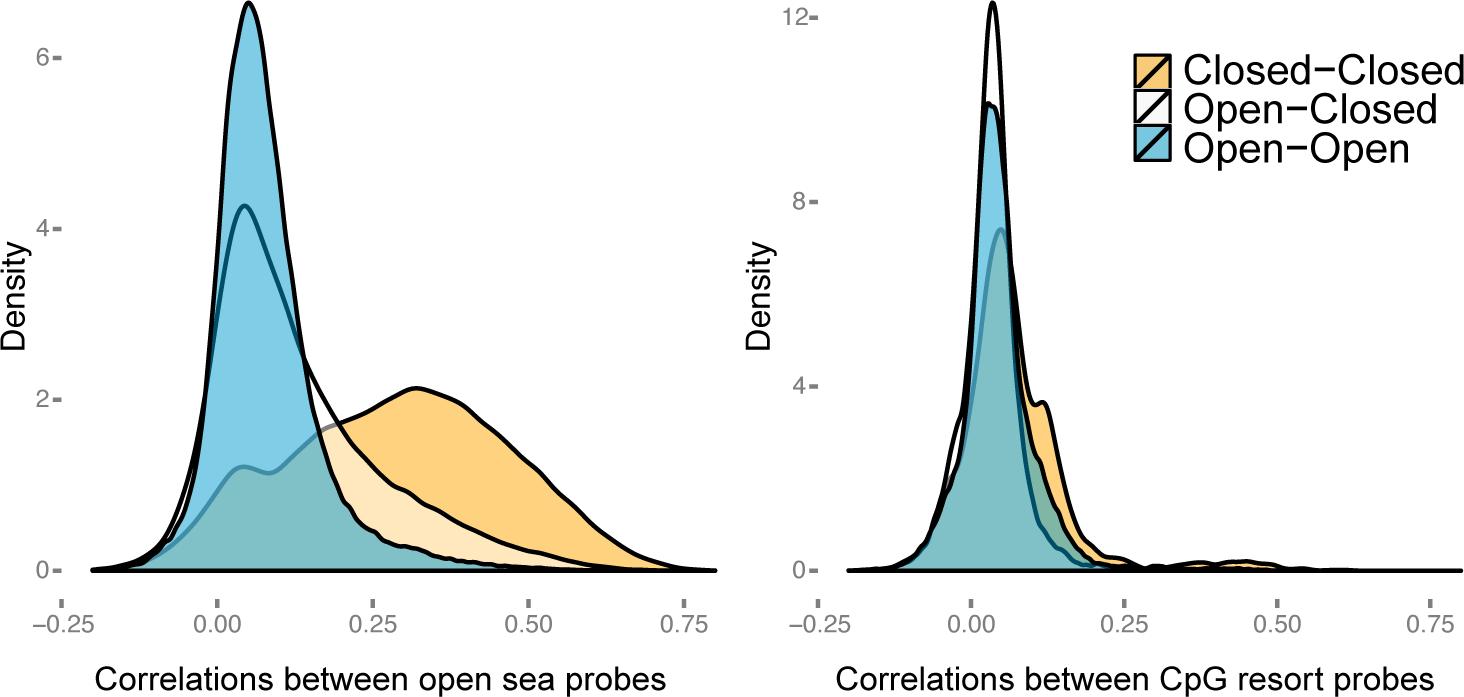
Densities of the correlations of the 450k methylation probes. Chromosome 14 was binned at resolution 100 kb and we display the binned, stratified correlations for the 450k-EBV dataset. Each plot shows one density curve for each type of interaction: between two bins in open compartmens, between two bins in closed compartments and between a bin in the open compartment and the closed compartment. (a) Binned correlations for open sea probes only. (b) Binned correlations for CpG resort probes only. Most correlations are around zero, except correlations between two open sea probes in the closed compartment. The open and closed compartments were defined using the EBV-2014 Hi-C dataset.

**Figure 8.**
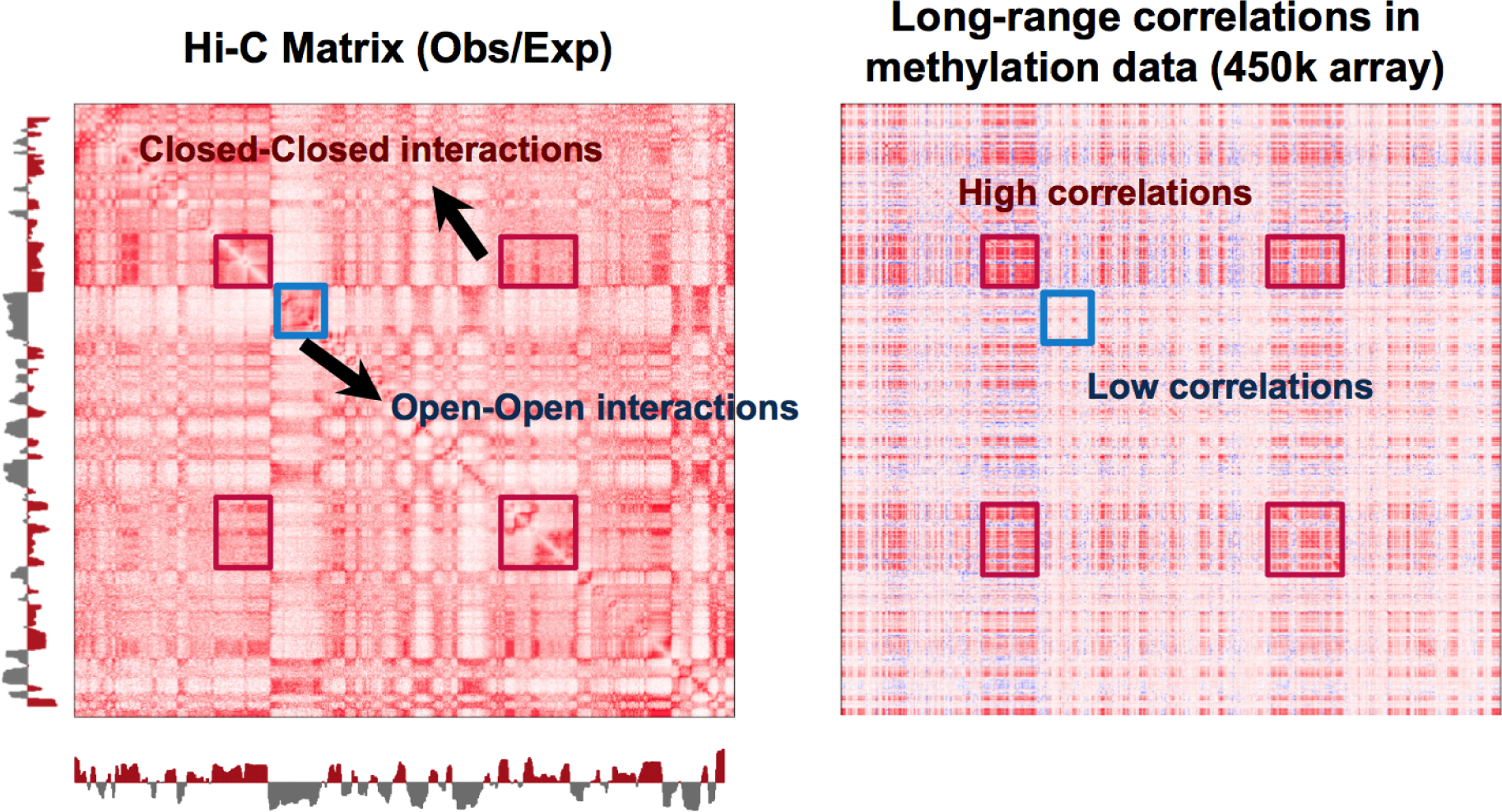
The relationship between a Hi-C contact matrix and a binned DNA methylation correlation matrix. Depicted are the observed-expected normalized genome contact matrix for the EBV-2014 Hi-C dataset together with the binned correlation matrix for the 450k-EBV dataset. Both matrices correspond to chromosome 14 at resolution 100kb. There is a relationship between A/B compartments in the Hi-C data and regions with low and high correlations.

The lack of decay of correlation with distance extends even to trans-chromosomal correlations, again with a clear difference between correlations within the open compartment and the closed compartment (Figure 9).

**Figure 9.**
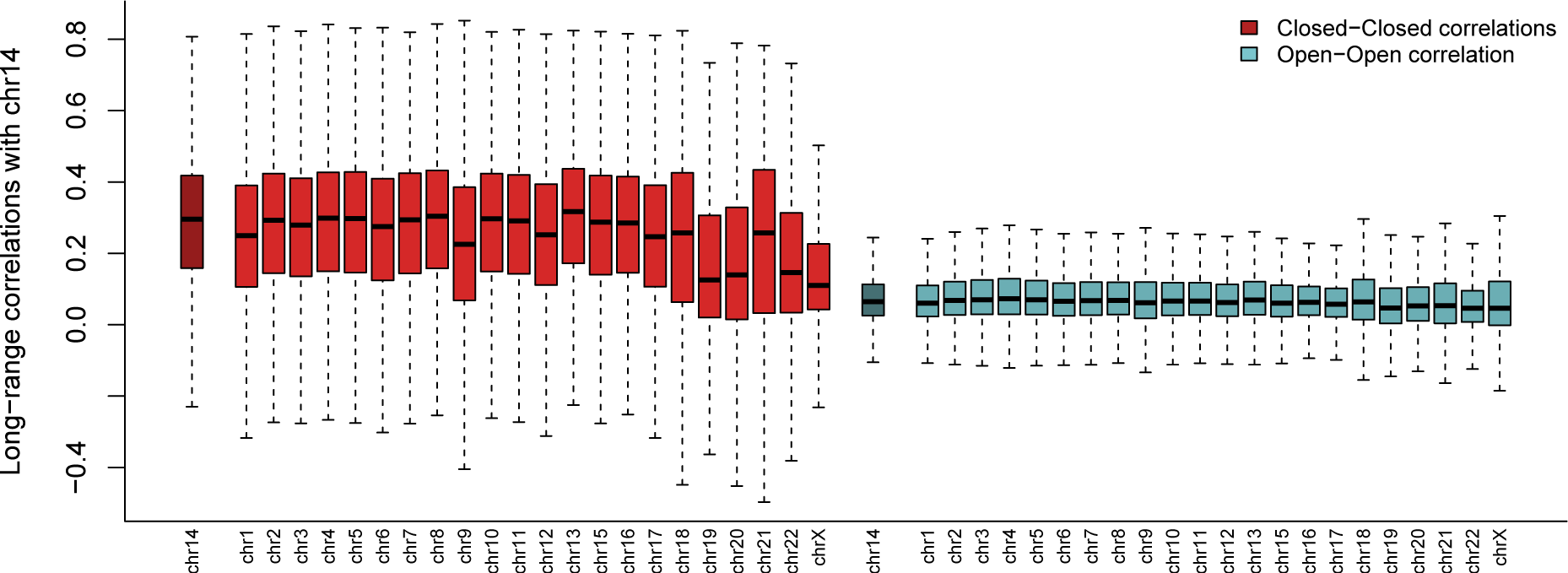
Between chromosome correlations of DNA methylation. Each boxplot shows the binned correlations between bins on chromosome 14 and bins on other chromosomes for the 450k-EBV dataset. The boxplots are stratified by whether the correlation is inside the open compartment or inside the closed compartment (open to close compartment correlations are not depicted). The open and closed compartments were defined using the EBV-2014 Hi-C dataset.

To understand what drives the correlation between closed compartments, we carefully examined the DNA methylation data in these genomic regions. Figure 10 shows a very surprising feature of the data, which explains the long-range correlations. In this figure, we have arbitrarily selected 10 samples and we plot their methylation levels across a small part of chromosome 14, with each sample having its own color. Data from both EBV-transformed lymphocytes and fibroblasts are depicted. While the same coloring scheme has been used for both cell types, there is no overlap between the samples assayed in the different experiments. The figure shows that the 10 samples have roughly the same ranking inside each region in the closed compartment. This illustrates a surprising genome-wide ranking between samples in the closed compartment.

**Figure 10.**
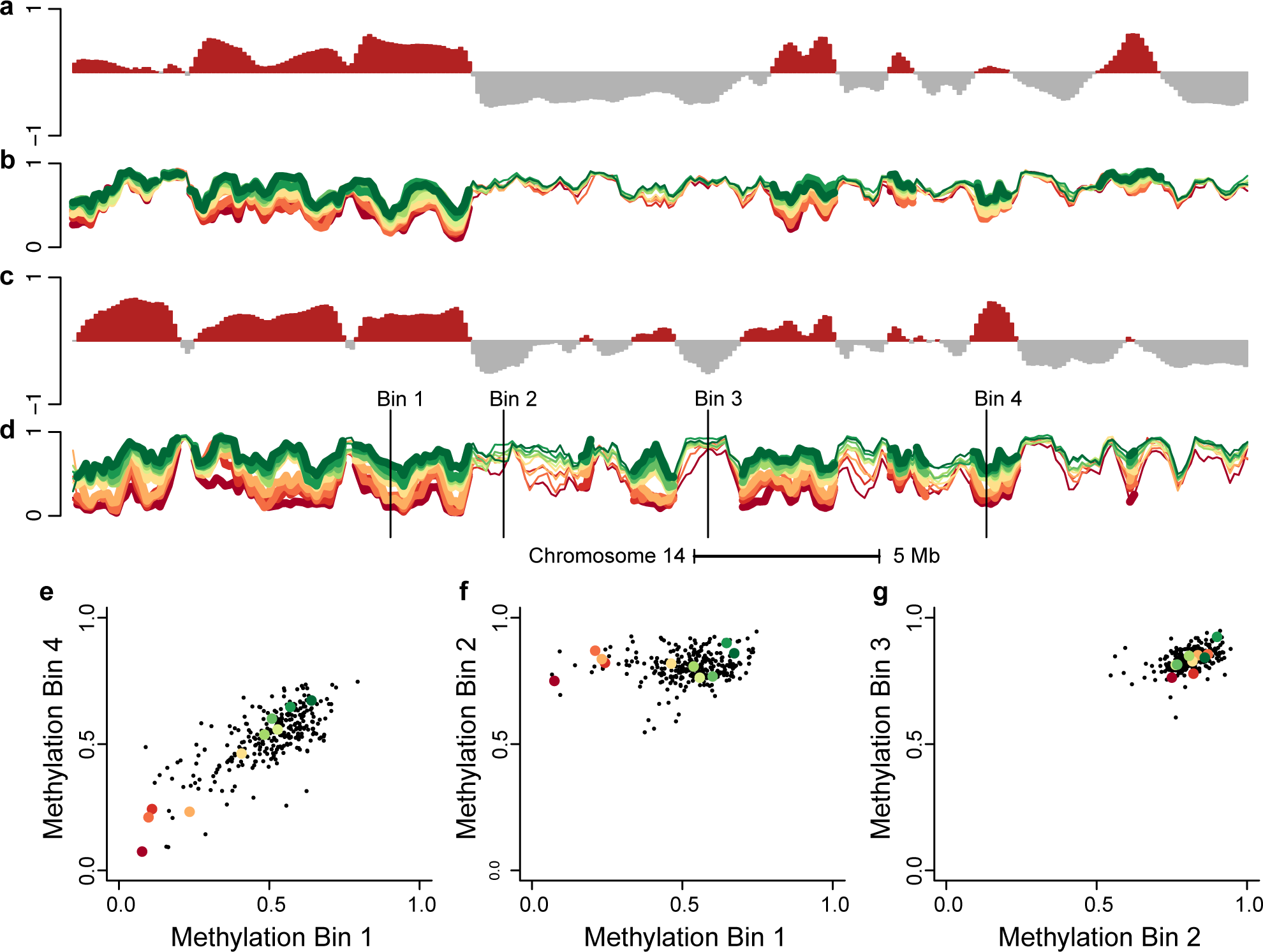
Comparison of the methylation levels and the Hi-C compartments signal for chromosome 14. The figure displays data from 36.4 to 69.8 Mb on chromosome 14 at 100kb resolution. (a) The first eigenvector from the IMR90-2014 Hi-C dataset. (b) Average methylation on the beta scale for 10 selected samples from the 450k-Fibroblast dataset; each sample is a line and divergent colors are used to distinguish the different levels of methylation in the different samples. (c) The first eigenvector from the EBV-2014 Hi-C data. (d) Like (b) but for 10 samples from the 450k-EBV dataset; the samples from the two datasets are unrelated. On (d) we depict 4 different bins; scatterplots between methylation values in different bins across all samples in the dataset are shown in (e-g) where (e) are two bins in the closed compartment, (g) are one bin in the open and one bin in the closed compartment and (g) two bins in the open compartment. The figure shows that samples have roughly the same ranking inside each closed compartment.

To gain more insights into whether this ranking was caused by technological artifacts or whether it reflects real differences between the biological replicates, we obtained data where the exact same HapMap samples were profiled in two different experiments using the Illumina 27k methylation array. This array design is concentrated around CpG islands, but we determined that 5,599 probes are part of the 450k array and annotated as open sea probes. For these probes, we determined which were part of the closed compartment and we computed the sample specific average methylation in this compartment, as a proxy for the observed ranking described above. In Figure 11a, we show that the correlation of these measurements between hybridization duplicates from the same experiment is high (92.7%). In Figure 11b we show that the these measurements replicate well between different experiments (correlation of 74.4%).

**Figure 11.**
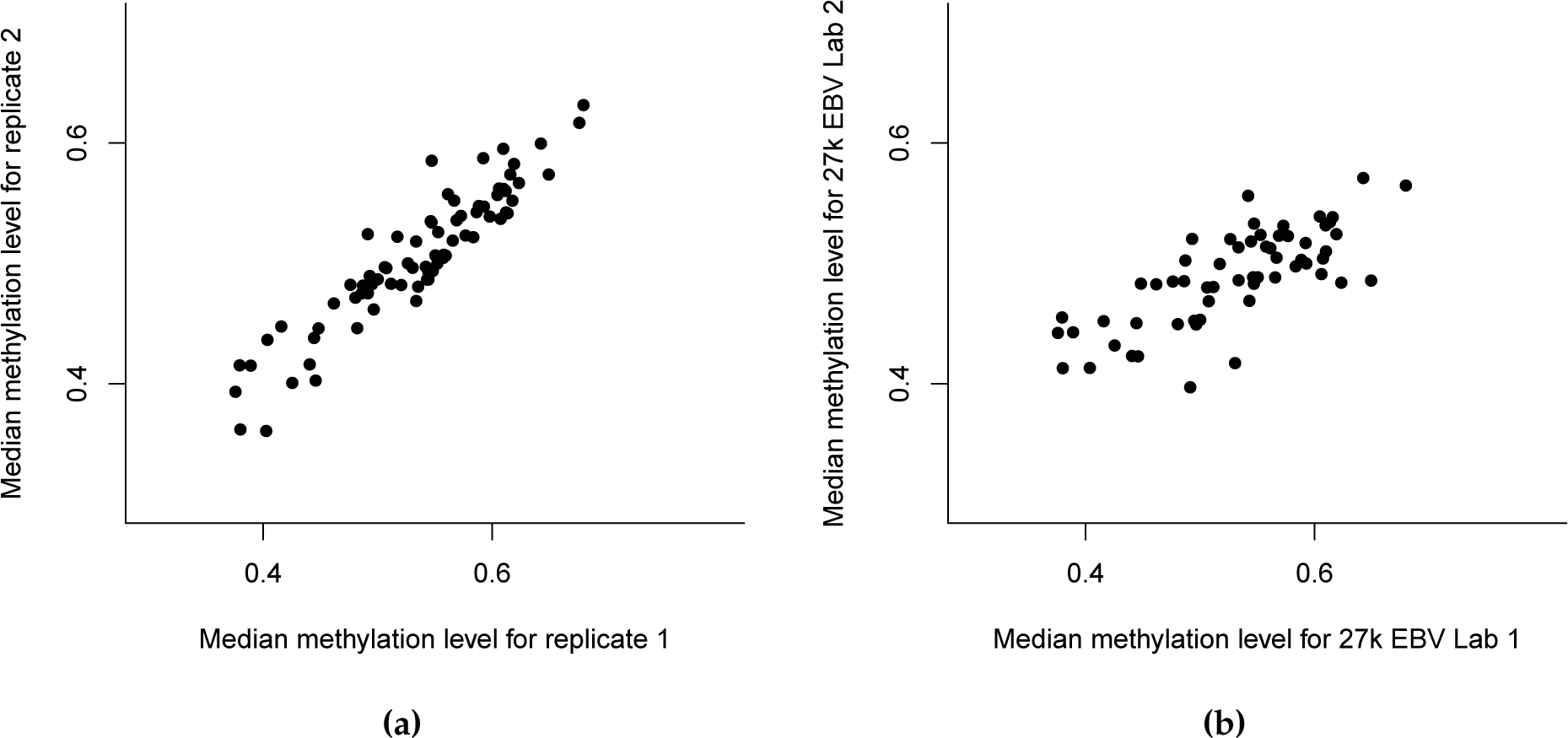
Sample ranking based on methylation levels in the closed compartments replicate across experiments. We computed the average methylation level of open sea probes in the closed compartment. The compartments were defined using the EBV-2014 Hi-C data. (a) Comparison between hybridization replicates from the 27k-London dataset. (b) Comparison between the same samples assayed in two different experiments, the 27k-Vancouver and the 27k-London experiments.

The striking global ranking between different samples using the open sea probes in the closed compartment could not be explained by the bisulfite conversion control probes, neither batch nor background noise. Neither the medians not the means were significantly associated with any of the different control probe types.

Finally, using the 27k data, we show that the eigenvector replicates between a 450k experiment and a 27k experiment using the same cell type (EBV), but different samples (correlation of 89%, see Figure 12). As a control, we compared to a 450k derived eigenvector for a different cell type (fibroblast) and observed weak correlation (40%). We note that the eigenvector derived from the 27k experiment is based on far fewer probes; we do not recommend using 27k data to estimate compartments. This result shows that the estimated genome compartments do not depend on the design of the microarray and suggests that our observations are common across methylation assays.

**Figure 12.**
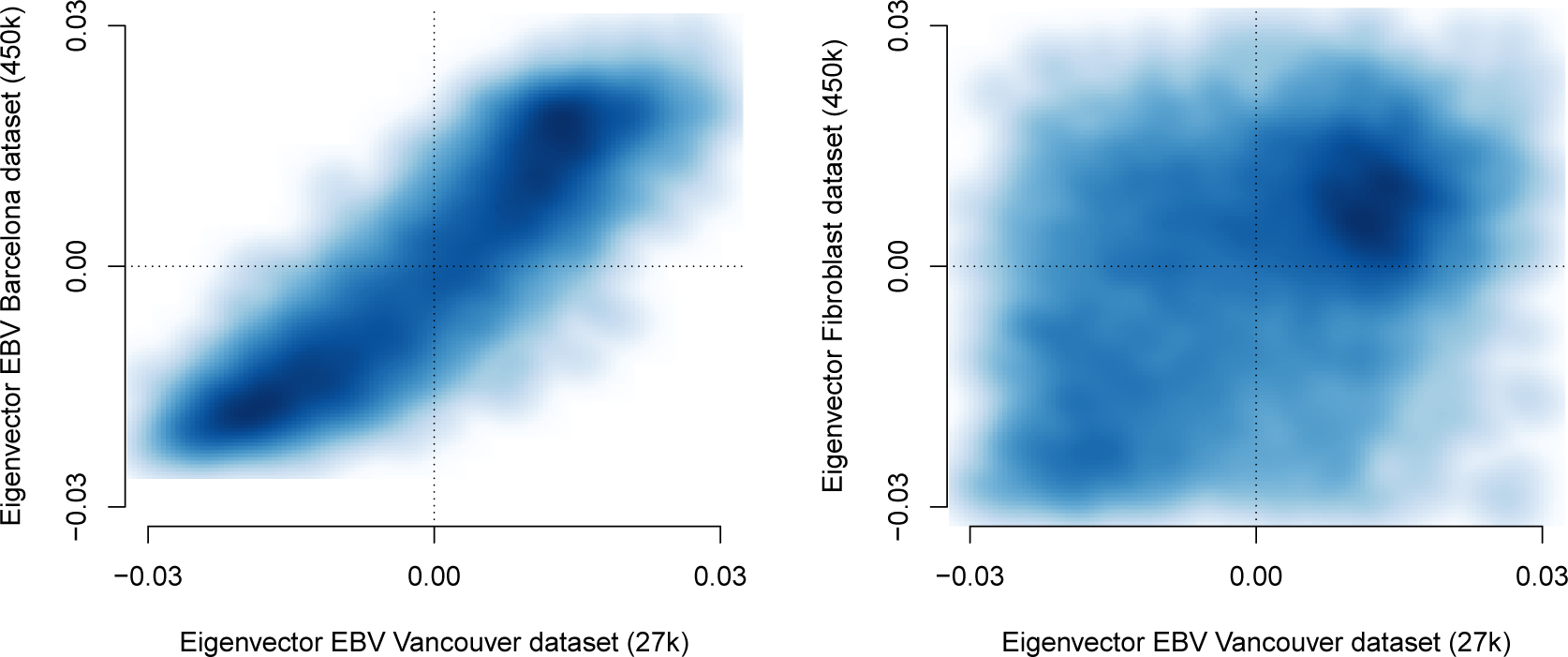
Validation of the 450k EBV eigenvector using a 27k dataset. The eigenvector for the 27k dataset was computed using the full (no binning) correlation matrix of all 5599 autosomal probes located in open sea which are commong between the two platforms. (a) A comparison between eigenvectors from the 27k-Vancouver dataset and the 450k-EBV dataset. The correlation between the two eigenvectors is 89.3%. (b) A comparison between eigenvectors from the 27k-Vancouver dataset and the 450k-Fibroblast dataset. The correlation is 42.1%, confirming the cell-type specificity of the methylation eigenvector.

### Notes on processing of the DNA methylation data

We have analyzed a wide variety of DNA methylation data both from the Illumina 450k and Illumina 27k microarrays. For each dataset, it varies which kind of data is publicly available (raw or processed). If possible, we have preferred to process the data ourselves starting from the Illumina IDAT files. However, for several datasets, we had to use the original authors preprocessing pipeline; see the Methods section for details.

We examined the impact of preprocessing methods on the estimated eigenvectors by using both functional normalization [Fortin et al., 2014b], quantile normalization adapted to the 450k array [Aryee et al., 2014] and raw (no) normalization; we did not find any substantial changes in the results. The agreement between the eigenvectors using the different preprocessing methods is greater than 94% and we note that the agreement with Hi-C data is best using functional normalization. This might be caused by the ability of functional normalization to preserve large differences in methylation between samples [Fortin et al., 2014b], which is what we observe in the closed compartment.

We examined the resolution of our approach using data from the 450k methylation array. As resolution increases, the number of bins with zero or few probes per bin increases. In Figure 13 we show the tradeoff between bins with zero probes and agreement with Hi-C data. This figure shows that a reasonable lower limit of resolution is 100kb. We note that the compartments estimated from Hi-C data do not change with increased resolution (Figure 2).

**Figure 13.**
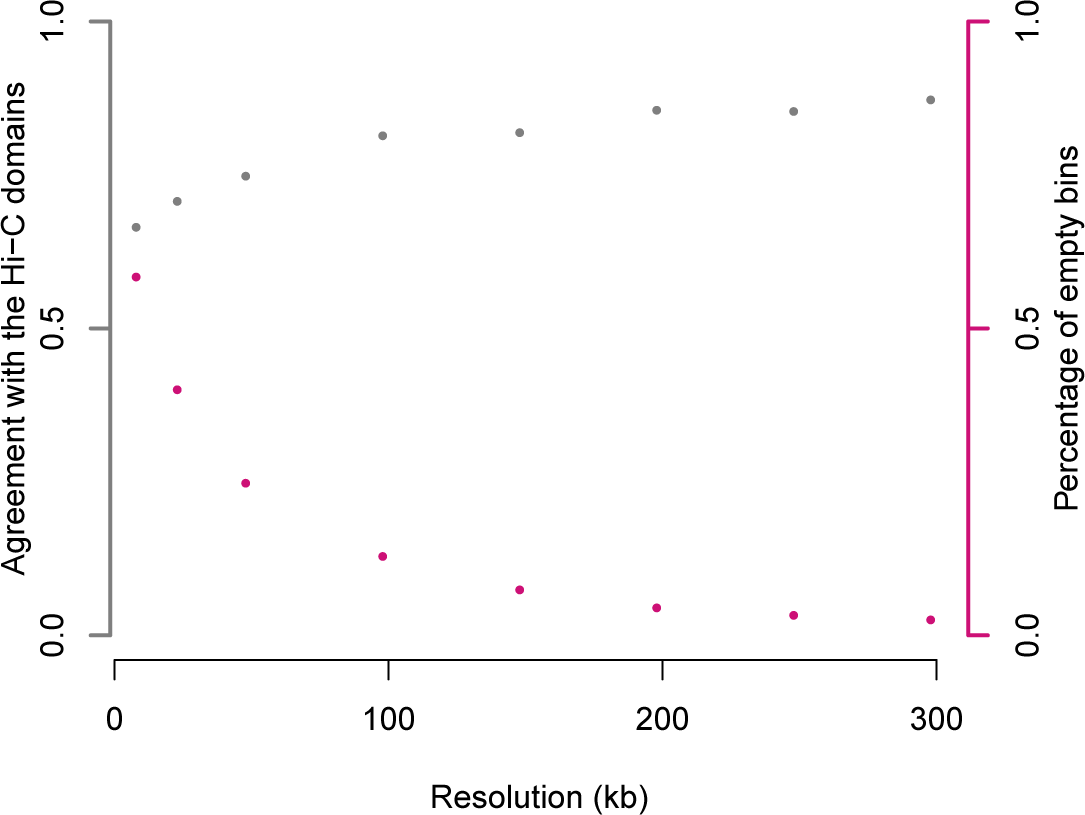
Performance of the A/B compartment prediction for different binning resolutions of the 450k array. The grey points depict the increase in agreement between compartments estimated using the 450k-EBV and EBV-2014 Hi-C datasets as binning resolutions coarsen. The pink points show the increase in empty bins as binning resolution is increased. We conclude that a resolution of 100kb is a good choice.

### An application to prostate cancer

We applied these methods to Illumina 450k data on prostate adenocarcinoma (PRAD) from The Cancer Genome Atlas (TCGA). Quality control shows both normal and cancer samples to be of good quality. Since the normal prostate samples represents uncultured primary samples, we confirmed that this dataset had the same information in its long-range correlation structure as established above (Figure 14; compare with Figure 10).

**Figure 14.**
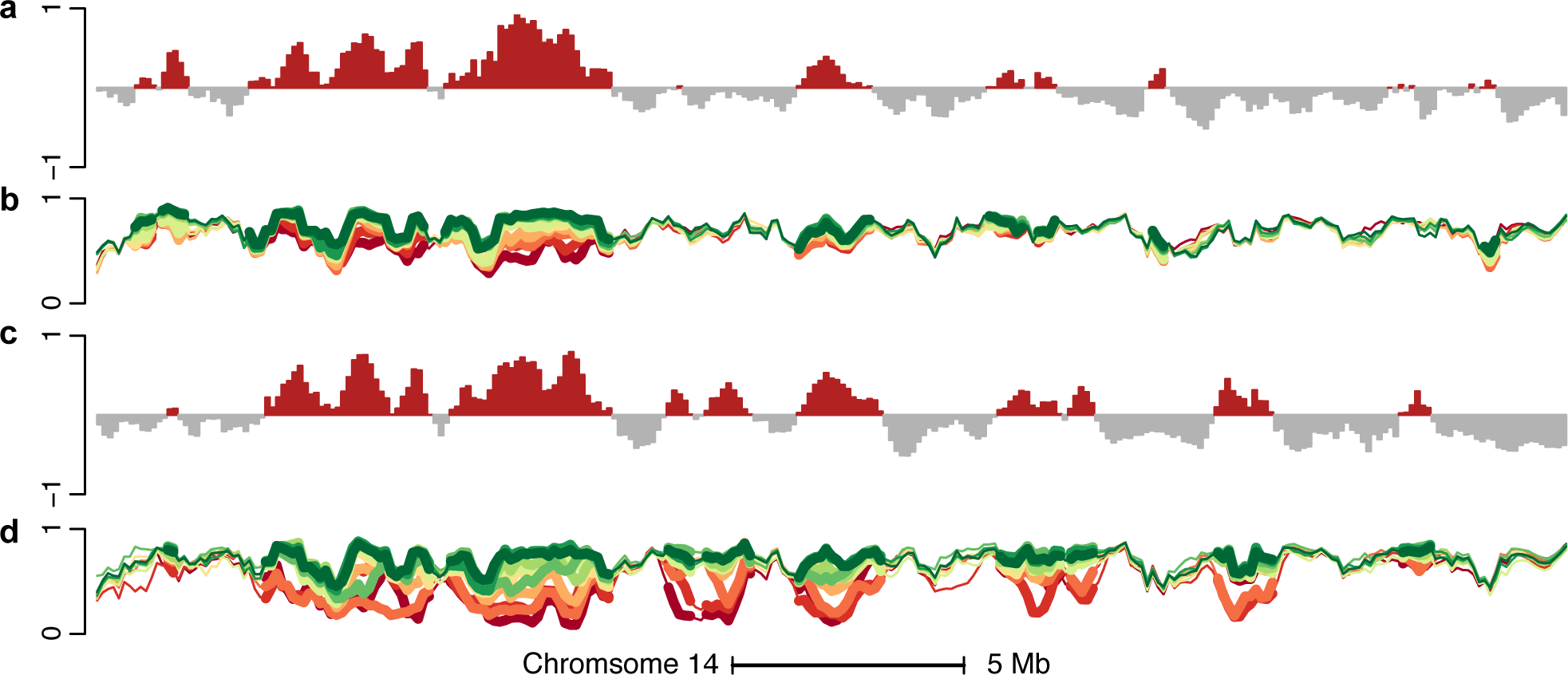
Comparison of the methylation levels and the HiC compartments signal for the TCGA PRAD datasets. Similar to Figure 10, but for the TCGA PRAD tumor/normal datasets. (a) The first eigenvector of the binned methylation correlation matrix for the TCGA PRAD normal dataset. (b) Average methylation signal on the beta scale for 10 selected samples for the TCGA PRAD normal dataset. (c) Like (a) but for the TCGA PRAD tumor dataset. (d) Like (b) but for the TCGA PRAD tumor dataset.

**Figure 15.**
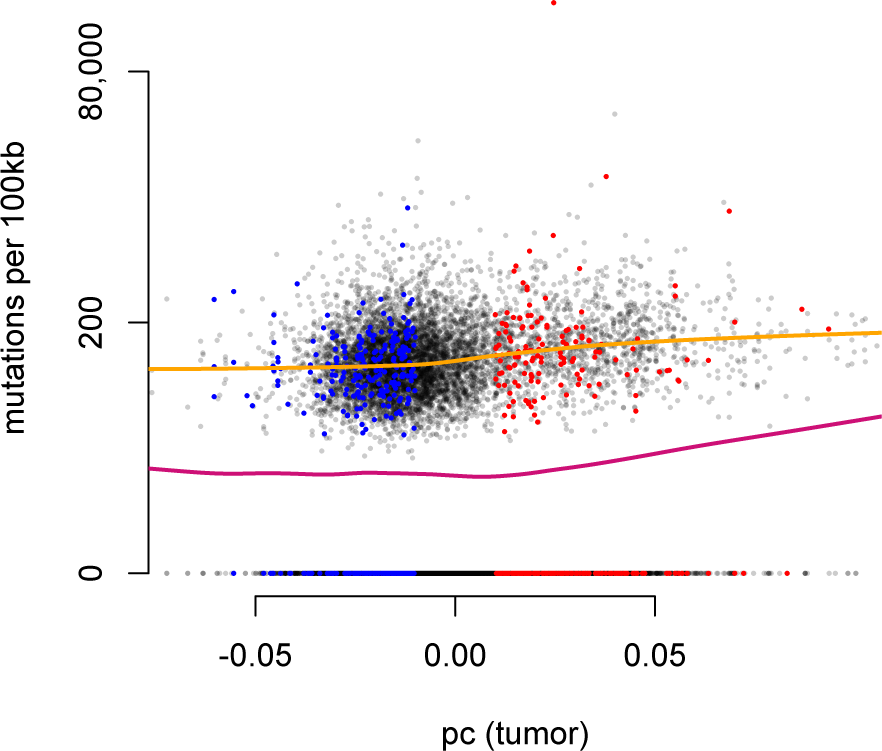
Relationship between A/B compartments and somatic mutation rate in prostate cancer. Somatic mutation rate for prostate cancer calculated using whole exome sequencing data from TCGA displayed against the first eigenvector of the 450k PRAD TCGA dataset. The y-axis uses the hyperbolic arcsine scale which is similar to the logarithm for values greater than one. A large number of genomic bins have a mutation rate of zero. The pink line is a lowess curve fitted to all the data and the orange line is a lowess curve fitted only to bins with a strictly positive mutation rate. We observe an increase in somatic mutation rate in the closed compartment, as expected. Colored points represent bins which confidently change compartments between normal samples and cancer samples, blue is closed to open and red is open to closed. A bin confidently changes compartment if its associated eigenvector value has a magnitude greater than 0.01 (but with different signs) in both datasets.

**Figure 16.**
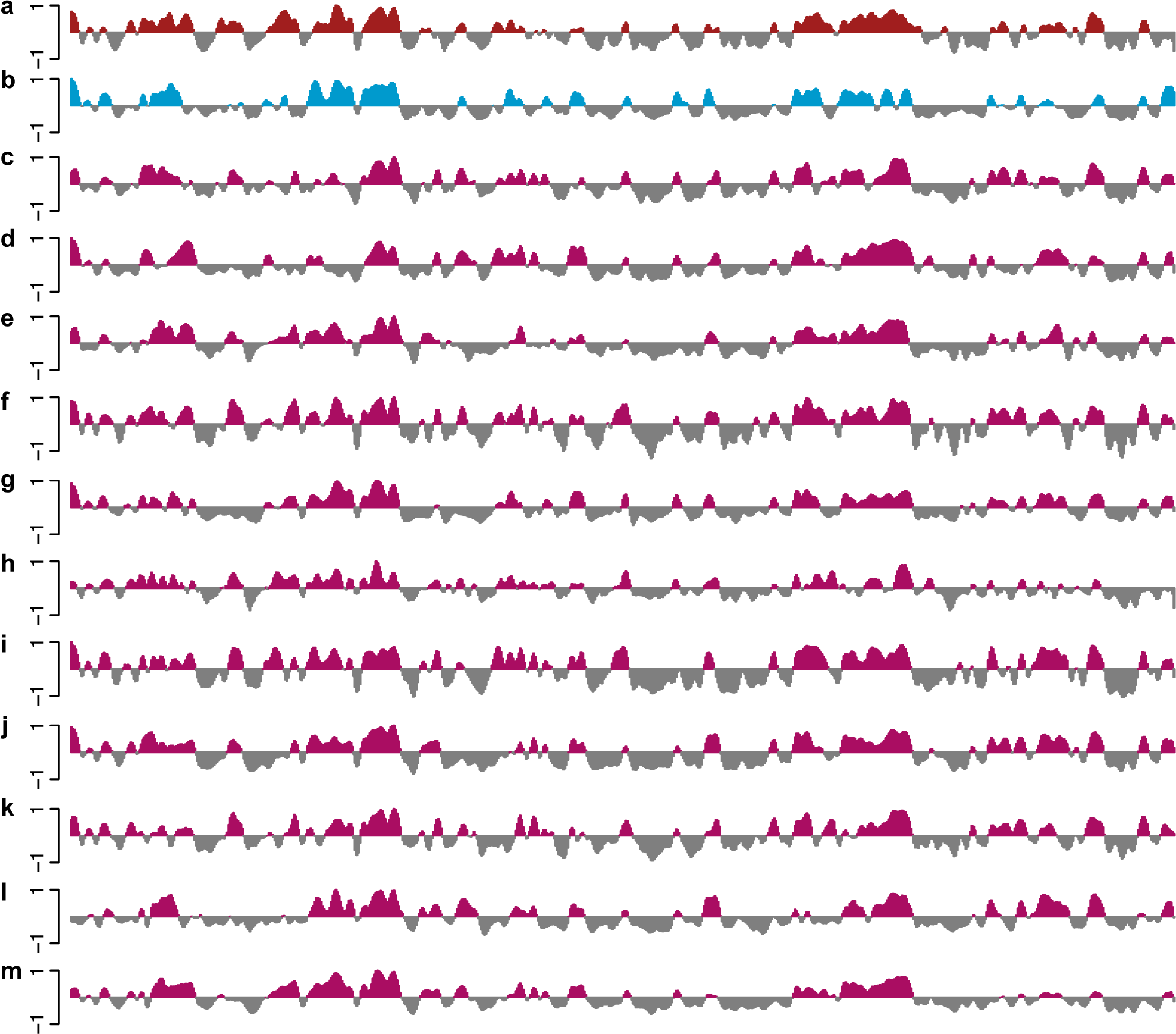
Estimated A/B compartments across several human cancers. The figure displays data on all of chromosome 14 at 100kb resolution. Each track represents the first eigenvector of the methylation correlation matrix for the corresponding dataset. The datasets depicted in (a) and (b) are the 450k-EBV and 450k-Fibroblast datasets. The datasets in (c-m) are cancer samples from TCGA for different cancers: (c) BLCA, (d) BRCA, (e) COAD, (f) HNSC, (g) KIRC, (h) KIRP, (i) LIHC, (j) LUAD, (k) LUSC, (l) PRAD, and (m) UCEC.

We obtained a list of curated somatic mutations from TCGA and used them to compute simple estimates of the mutation rate in each 100kb bin of the genome. Since the list of somatic mutations was obtained using whole-exome sequencing (WXS), we obtained the relevant list of capture regions and used this list to compute mutation rates. We compared the somatic mutation rate to the eigenvector estimating the open and closed compartments and found the mutation rate to be elevated in the closed compartment. This confirms previous observations about the relationship between mutation rates and open and closed chromatin [Makova and Hardison, 2015], including cancer [Schuster-Böckler and Lehner, 2012, Polak et al., 2015]. Of particular interest was the mutation rate in genomic regions belonging to different compartments in normal and tumor samples. Table 2 shows these mutation rates computed using bins where the associated eigenvector value has a magnitude greater than 0.01; this was done to discard bins where the compartment association could be considered ambiguous. The table shows that regions of the genome changing from open to closed compartment in tumors have a similar mutation rate to regions which are in the closed compartment in both tumor and normals. Said differently: changes in compartments are associated with changes in somatic mutation rate. To our knowledge, this is the first time a cancer-specific map of open and closed compartments based on primary samples have been derived; existing analyses depends on chromatin assays performed in ENCODE and Epigenomics Roadmap samples [Schuster-Böckler and Lehner, 2012, Polak et al., 2015].

**Table 2.** Mutation rate per 100kb of somatic mutations in PRAD stratified by compartment.

|  |  | Normal |  |
| --- | --- | --- | --- |
|  |  | Open | Closed |
| Tumor | Open | 54.1 | 58.0 |
|  | Closed | 83.9 | 97.2 |

### Compartments across human cancers

Using the method we have developed in this manuscript, it is straightforward to estimate A/B compartments across a wide variety of human cancers using data from TCGA. Figure 16 displays the smoothed first eigenvectors for chromosome 14 at 100kb resolution for eleven different cancers. Regions of similarity and differences are readily observed. We emphasize that TCGA does not include assays measuring chromatin accessibility such as DNase or various histone modifications. The extent to which these differences are associated with functional differences between these cancers is left for future work.

### Compartment prediction using DNase hypersensitivity data

Lieberman-Aiden et al. [2009] established a connection between A/B compartments and DNAse data, mostly illustrated by selected loci. Based on these results we examined the degree to which we can predict A/B compartments using DNase hypersensitivity data. This data, while widely available from resources such as ENCODE, does not encompass the wide variety of primary samples as the Illumina 450k methylation array.

We obtained DNase-seq data on 70 samples [Degner et al., 2012] from EBV-transformed lymphocytes from the HapMap project, as well as 4 experiments on the IMR90 cell line performed as part of the Roadmap Epigenomics project [Bernstein et al., 2010]. We computed coverage vectors for each sample and adjusted them by library size. For each sample, we computed the signal in each 100kb genomic bin and averaged this signal across samples. The resulting mean signal is highly skewed towards positive values in the open compartment, and we therefore centered the signal by the median. The median was chosen as this has the best compartment agreement with Hi-C data. Figure 17 shows the result of this procedure, slightly modified for display purposes (the sign was changed to let high values be associated with closed compartment; additionally very low values were thresholded). A good visual agreement is observed for both cell types; the correlation between Hi-C and the average DNase signal on chromosome 14 is 72% for EBV and 76% for IMR90 with a compartment agreement of 83% for EBV and 86% for IMR90.

**Figure 17.**
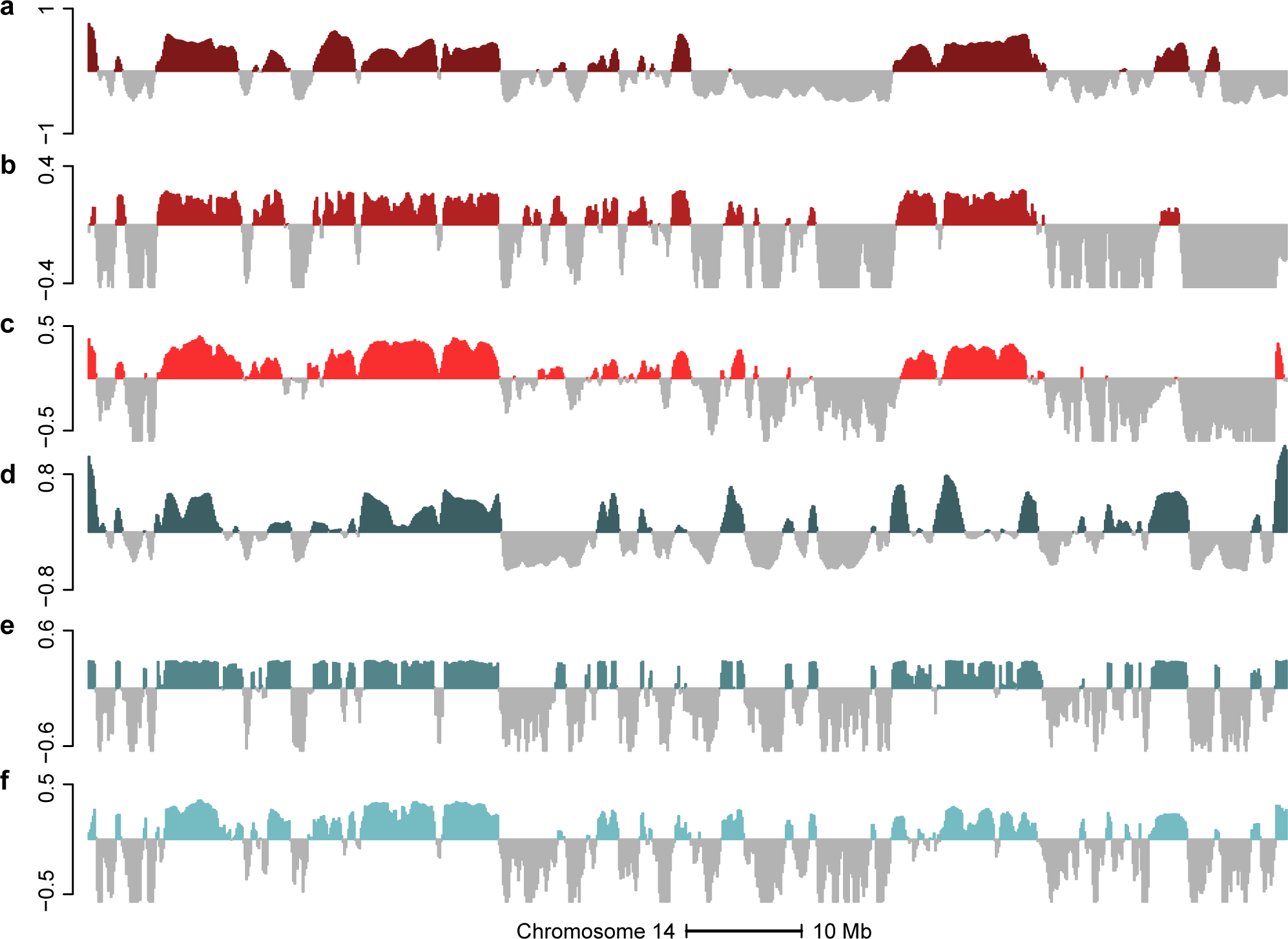
DNase data can predict A/B compartments revealed by Hi-C. The figure displays data on all of chromosome 14 at 100kb resolution. (a) The first eigenvector of the EBV-2014 Hi-C dataset. (b) The smoothed first eigenvector of the correlation matrix of the binned DNase-EBV dataset after median centering. (c) Average DNase signal across samples after binning and median subtraction. The sign of the signal was reversed for display purpose. (d) The first eigenvector of the IMR90-2014 Hi-C dataset. (e) The smoothed first eigenvector of the correlation matrix of the binned DNase-IMR90 dataset after median centering. (f) Average DNase signal across samples after binning after median subtraction. The sign of the signal was reversed for display purpose. Both the average signal and correlation eigenvector are highly predictive of the Hi-C compartments for both cell types.

Inspired by the success of considering long-range correlations for the 450k data, we computed a DNase correlation matrix by computing the Pearson correlation matrix of the binned DNase signal; in contrast to the 450k data, we did not bin the correlation matrix as the signal matrix was already binned. The first eigenvector of this correlation matrix is highly skewed; we centered it by its median. Figure 17 shows the result of this procedure. We obtained a correlation between this centered eigenvector and the Hi-C eigenvector of 78% for EBV and 76% for IMR90 and a compartment agreement of 86% for EBV and 83% for IMR90. These results are comparable to what we obtain using the average DNase signal. It might be notable that the correlation based method works better for the EBV data which contains biological replicates and worse for the IMR90 dataset which is based on growth replicates of the same cell line.

To examine why the correlation based approach works for DNase data, we performed the same investigation as for the 450k datasets. In Figure 18 we show the distribution of correlations stratified by compartment type. As for the DNA methylation data, the DNase data has high positive correlations between bins in the closed compartment, although the correlations in the DNase data are much higher. For the DNA methylation data, correlations were close to zero between loci when at least one loci was in the open compartment. In contrast to this, the DNase data show an almost uniform distribution of correlation values when one of the two loci are in the open compartment.

**Figure 18.**
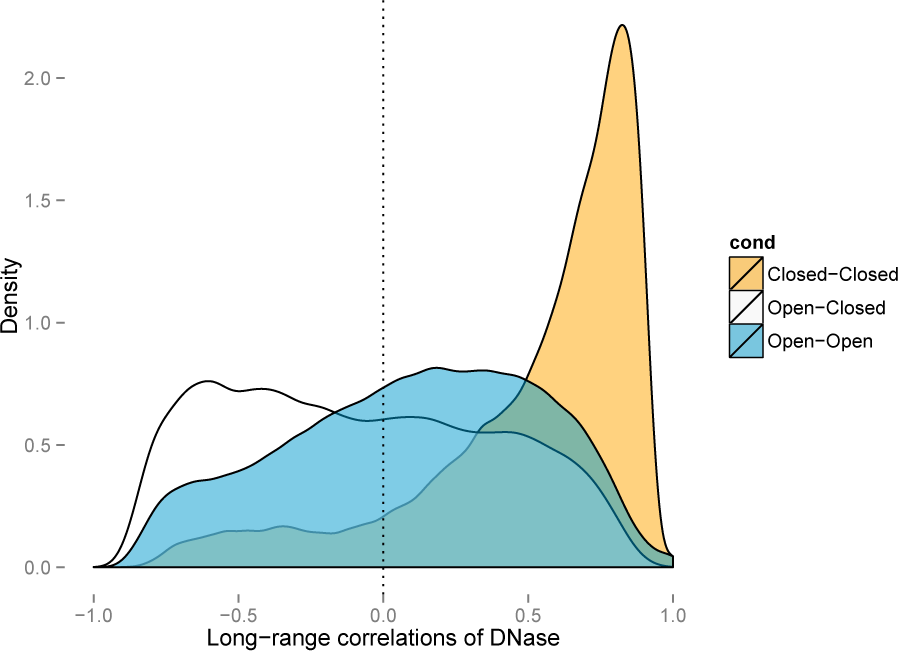
Densities of the correlations of DNase data. Chromosome 14 was binned at resolution 100kb. Depicted is the correlations of this data for the DNase-EBV dataset, stratified by compartment type. The open and closed compartments were defined using the EBV-2014 Hi-C dataset. This figure is similar to Figure 7.

Figure 19 suggests that, like DNA methylation, the DNase signal is ranked in the same way between samples in every region part of the closed compartment. There is a tendency for the ranking to be reversed in the open compartment.

**Figure 19.**
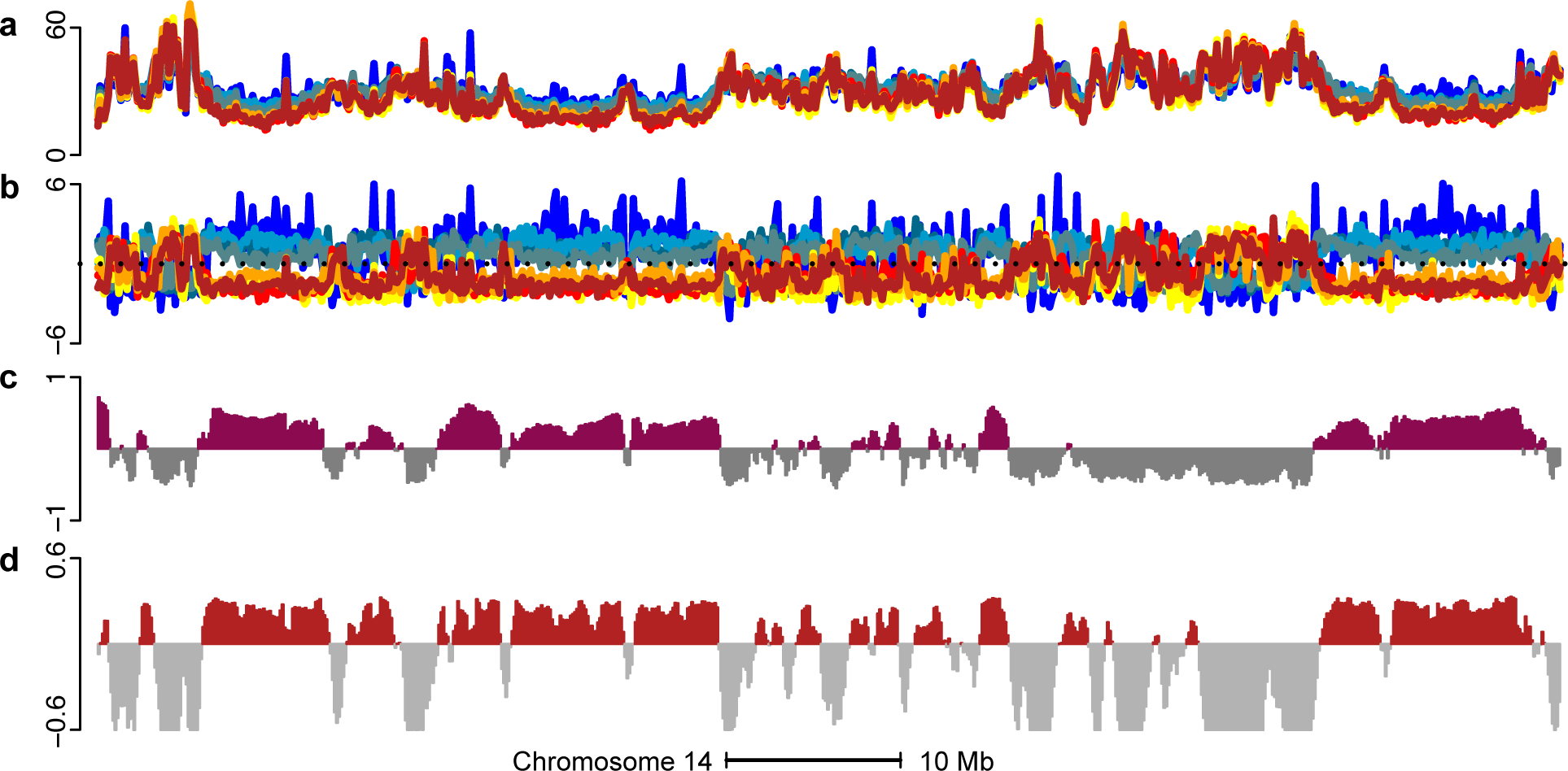
DNase samples have consistent ranking within closed compartments. The figure displays data from 20.2 to 90.1 Mb on chromosome 14 at 100kb resolution.(a) DNase signal for a subset of samples (scaled by square root for display purpose) (b) Same as (a), but centered at each position to look at variation of the signals with respect to the mean signal (original scale) (c) First eigenvector of the Hi-C EBV 2014 dataset (d) First eigenvector of the DNase-EBV correlation matrix. The signal below is 0 is truncated for display purpose.

### Compartment prediction using single-cell epigenetic data

Experimental techniques for measuring epigenetics in a single cell are in rapid development. We have applied our methods to data from the few genomewide, single-cell epigenetic experiments available. This includes data both on chromatin accessibility [Cusanovich et al., 2015] and DNA methylation [Smallwood et al., 2014].

Chromatin accessibility is measured by a single cell variant of an assay called assay for transposase-accessible chromatin (ATAC) sequencing [Buenrostro et al., 2013], which generates data similar to DNA hypersensitivity. From Cusanovich et al. [2015] data is available on mixtures of two cell lines, GM12878 and HL60, but not on pure samples of one cell type. First, we developed a simple method for assigning single cells from this mixture to one of the two known cell lines, based on average accessibility of known, cell-type specific hypersensitive sites; this is a much simpler method than what is suggested in Cusanovich et al. [2015]. Using our method, we observe two distinct clusters of cells, and it was easy to assign most cells unambiguously to a cell type using an arbitrary but seemingly sensible cutoff (Methods, Figure 20). This yielded data on 2679 cells from the GM12878 cell line from one experiment. We next applied our correlation based approach to this data; now the correlation is between single cells within the same cell line. Furthermore, the data consists of accessibility quantified over 195882 hypersensitive sites the original authors derived from ENCODE data, with the accessibility of each site being a value of 0, 1, or 2. We summarize this data in 100kb bins (see Methods), not unlike our treatment of bulk DNase-seq data. On chromosome 14 we observe a correlation of 0.83 and a compartment agreement of 81% between the first eigenvector of this data and the eigenvector from EBV-2014 Hi-C data (Figure 21).

**Figure 20.**
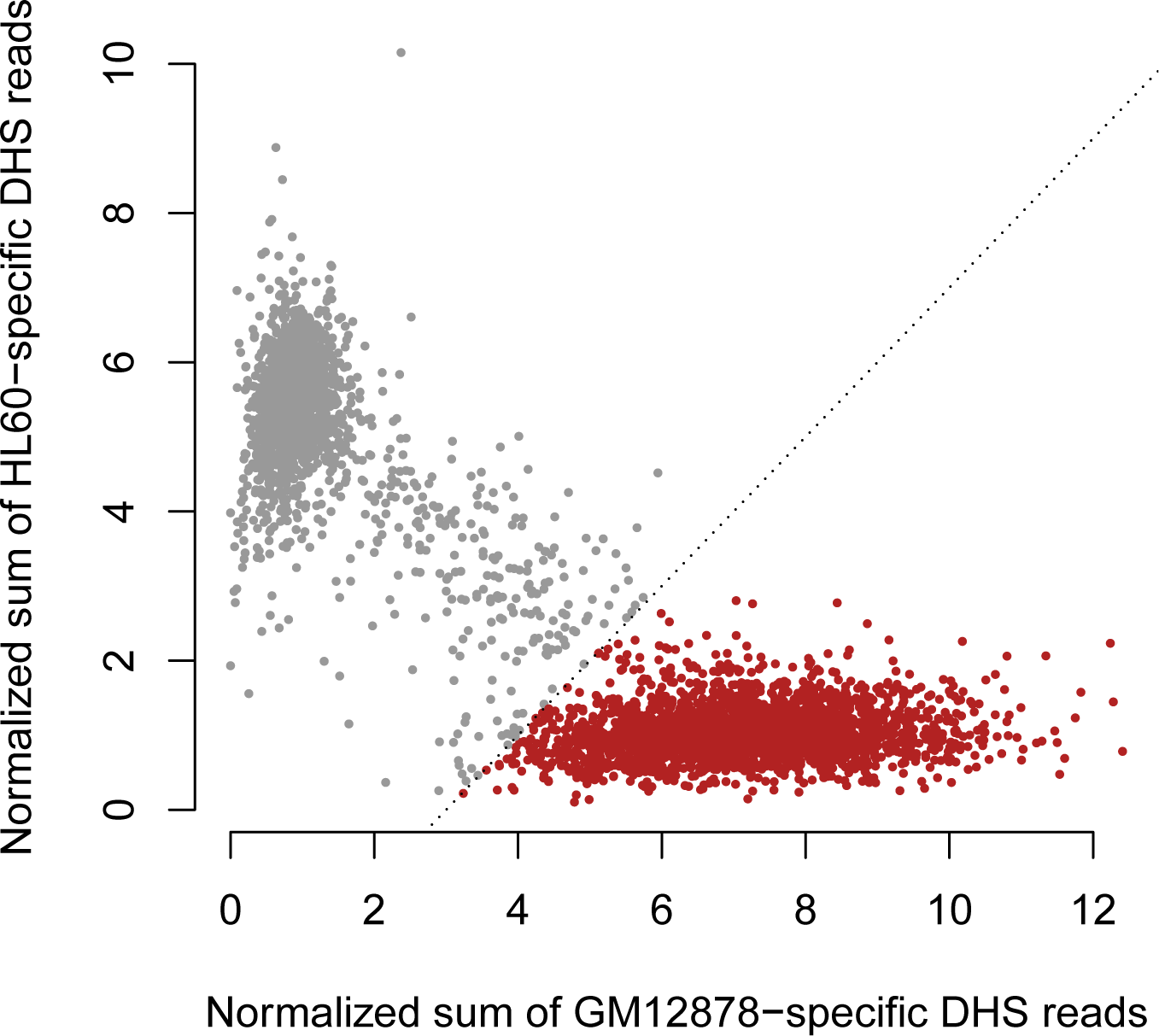
Cell type classification of single cell ATAC-seq data. Depicted is data from a single replicate of a mixture of the GM12878 and HL60 cell lines described in Cusanovich et al. [2015]. ENCODE DNAse-seq data was used to define hypersensitive sites (DHS) specific to these two cell lines. For each of these two sets of sites we computed the average number of ATAC-seq reads normalized by the total number of reads mapped to known DHS sites. The figures shows two distinct clusters; we arbitrarily selected the line *y* = *x* – 3 to delineate cells from the GM12878 cell line (red points).

**Figure 21.**
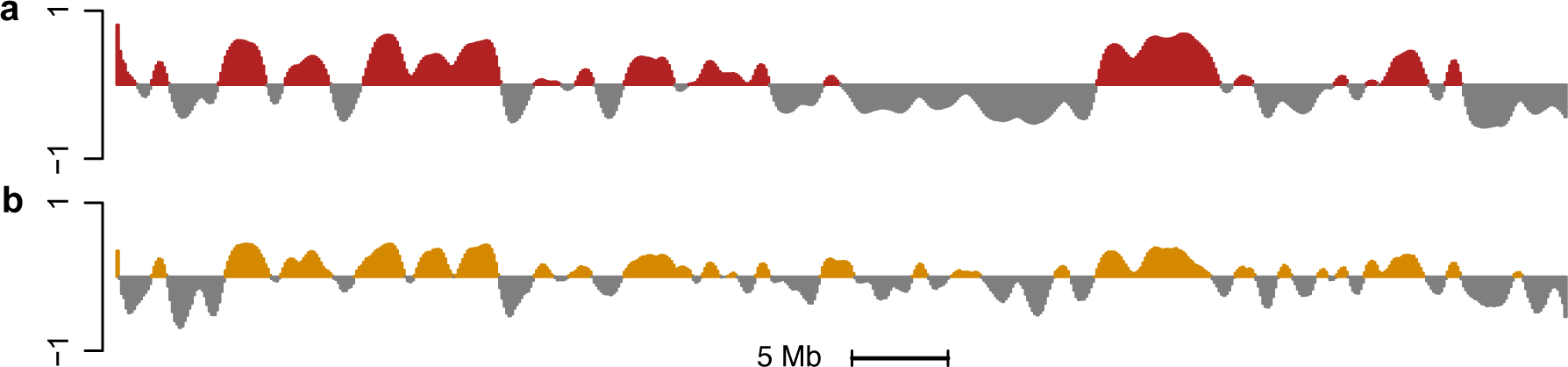
Estimated compartments from single-cell ATAC-seq data. Depicted is data from EBV transformed lymphocytes on chromosome 14 at a resolution of 100kb. (a) Estimated compartments using the EBV-2014 Hi-C data. (b) Estimated compartments using single-cell ATAC seq data across 2679 cells from GM12878.

Single cell DNA methylation can be measured using a form of whole-genome bisulfite sequencing (WGBS) as described in Smallwood et al. [2014]. Due to technical limitations of the assay, the number of assayed cells is small. We have data on 20 individual mouse embryonic stem cells (mESC) cultured in serum conditions, with corresponding Hi-C data from a different source [Dixon et al., 2012]. We generated a binned methylation matrix by averaging methylation values for open sea CpGs and discarded bins with little or no data (see Methods). We next applied our correlation based approach to this data, computing a correlation matrix across these 20 cells. On mouse chromosome 12 we observed a correlation of 0.61 and a domain agreement of 81%, using existing Hi-C data on the mouse embryonic stem cell line J1 [Dixon et al., 2012] (Figure 22). An analysis of the pattern of correlation between loci in open and closed compartments show some difference between the two distributions (Figure 23). In contrast to what we observe for 450k data, loci in the open domain are still substantially positively correlated. We note that Smallwood et al. [2014] show substantial between-cell heterogeneity in genomewide methylation across these 20 cells, depicted in Figure 23. However, this heterogenity of genomewide methylation was not observed for mouse ovulated metaphase II oocytes (MII) cells (Figure 23); the correlation distribution is substantially different for this dataset (Figure 23) and the first eigenvector of the correlation matrix only explains 19% of the variance, in contrast to 99% of the variance explained for mESC cells. The first eigenvector is depicted in Figure 22. We do not have Hi-C data available for this cell type, but we are doubtful that the first eigenvector accurately reflects the A/B compartments in this cell type.

**Figure 22.**
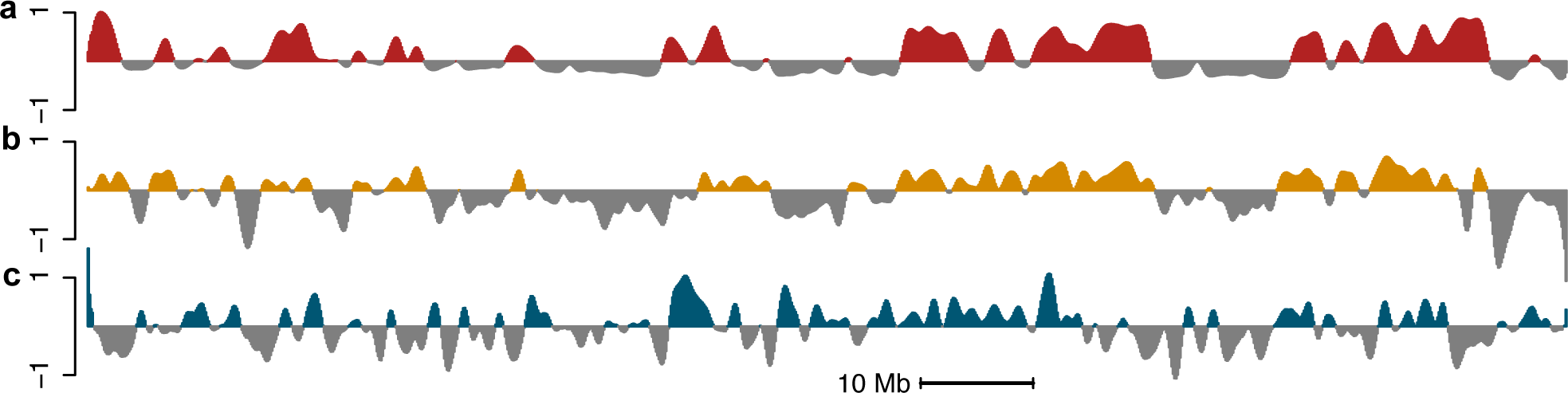
Estimated compartments from single-cell WGBS data. Depicted is data from mouse embryonic stem cells, chromosome 12 at a resolution of 100kb. (a) Estimated compartments using the mESC-2012 Hi-C data. (b) Estimated compartments using single-cell whole-genome bisulfite sequencing data from 20 mESC cells grown on serum. (c) The first eigenvector of a correlation matrix obtained using single-cell whole-genome bisulfite sequencing data from 12 ovulated metaphase II oocytes cells.

**Figure 23.**
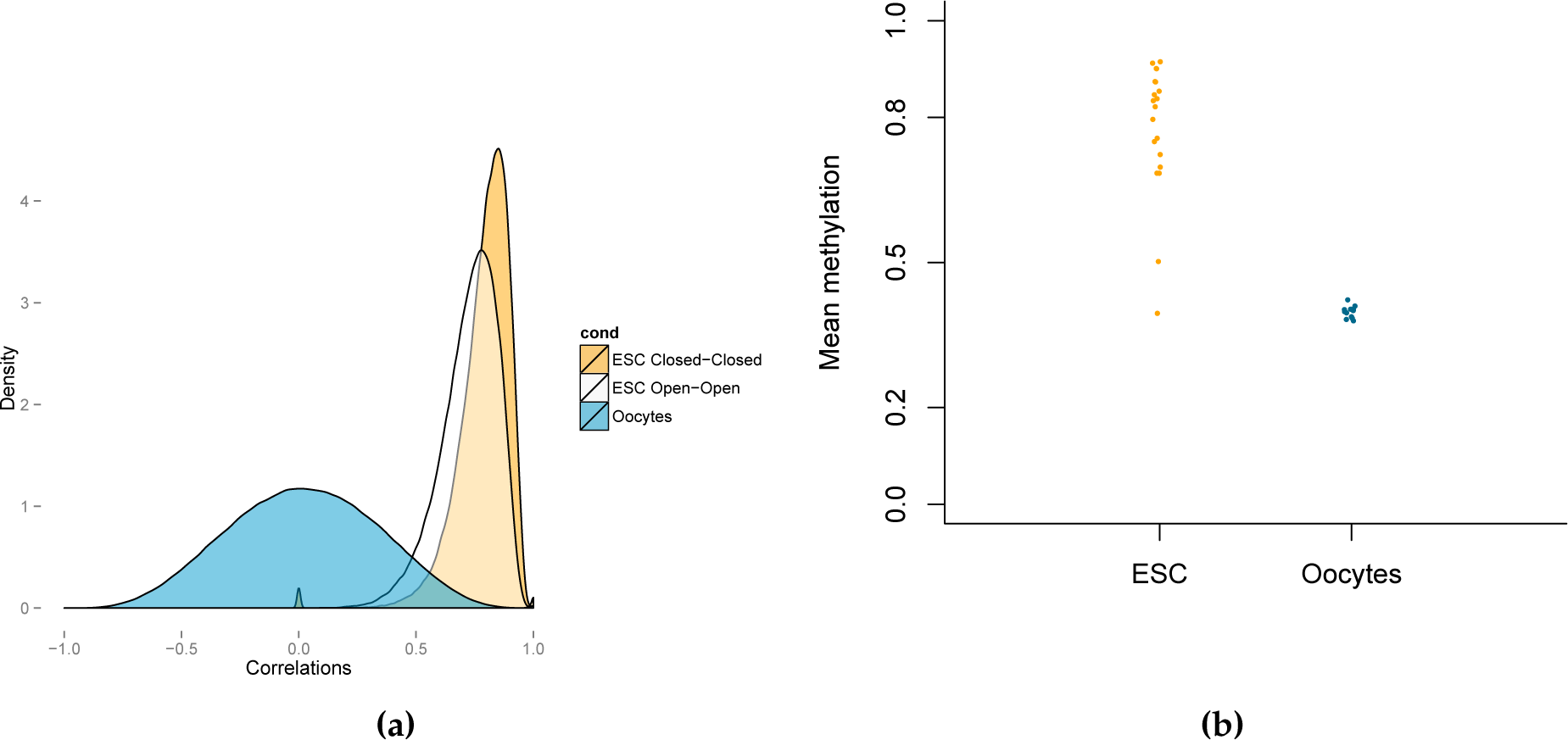
Structure of single-cell WGBS data. (a) Density of correlations for data on mESC and MII cells; compartments are estimated using the mESC-2012 Hi-C data. The two cell types have very different patterns. (b) Genome-wide methylation for 20 mouse ESC cells and 12 ovulated metaphase II oocytes (MII) cells. Substantial heterogeneity is observed for the former but not the later.

## Discussion

In this work, we show how to estimate A/B compartments using long-range correlations of epigenetic data. We have comprehensively evaluated the use of data from the Illumina 450k DNA methylation microarray for this purpose; such data is widely available on many primary cell types. Using data from this platform, we can reliably estimate A/B compartments in different cell types, as well as changes between cell types.

This result is possible because of the structure of long-range correlations in this type of data. Specifically, we found that correlations are high between two loci both in the closed compartment and low otherwise, and does not decay with distance between loci. This result only holds true for array probes measuring CpGs located more than 4kb from CpG islands, so-called open sea probes. This high correlation is the consequence of a surprising ranking of DNA methylation in different samples across all regions belonging to the closed compartment. We have replicated this result in an independent experiment using the Illumina 27k DNA methylation microarray.

We have furthermore established that A/B compartments can be estimated using data from DNase hypersensitivity sequencing. This can be done in two ways: first by simply computing the average DNase signal in a genomic region, and second by considering long-range correlations in the data, like for 450k array data. Again, we exploited the structure of long range correlations in this type of epigenetic data and, like the case for DNA methylation data, we found that correlations between loci both in the closed compartment are high, whereas correlations between other loci are approximately uniformly distributed. Again, this correlation is caused by a ranking of DNase signal in different samples across all regions belonging to the closed compartment. Surprisingly, our method works both for biological replicates (EBV transformed lymphocytes) but also on technical between-lab replicates of the same cell line (IMR90).

Finally we have established that our method works on single cell epigenetic data, including single cell ATAC-seq and single cell WGBS. These experimental techniques are in their infancy; it is likely that additional data will allow us to tune aspects of our method to this type of data. Now, the correlation is between single cells as opposed to biological replicates of bulk cells. This potentially allows our method to be used on rare types of cells.

Our approach is based on computing the first eigenvector of the (possibly binned) correlation matrix. It is well known that this eigenvector is equal to the first left singular vector from the singular value decomposition of the data matrix. The right singular vector of this matrix is in turn equal to the first eigenvector of the sample correlation matrix; also called the first principal component. This vector has been shown to carry fundamental information about batch effects [Leek et al., 2010]. Because of this relationship, we are concerned that our method might fail when applied to experiments that are heavily affected by batch effects; we recommend careful quality control of this issue before analysis.

The reason our method works is because of a surprising, consistent ranking of different samples across all regions belonging to the closed compartment (and only the closed compartment). By comparison with additional 27k methylation array experiments, we have shown that this ranking is not a technical artifact caused by (for example) hybridization conditions. In recent work, we studied colon cancer and EBV transformation of lymphocytes using WGBS [Hansen et al., 2011, 2014]. Using WGBS, it is easy to estimate the average methylation level across all CpGs in the genome; we call this global methylation. In these two systems, we observed global hypomethylation as well as an increased variation in global methylation levels in colon cancer and EBV-transformed lymphocytes when compared to normal matched samples from the same person. However, we saw minimal variation in global methylation between 3 normal samples in both systems. This work might explain why EBV-transformed samples from different people show a consistent ranking genomewide. But it does not explain why the same observation is made in fibroblasts and normal primary prostate (however, the later could be affected by contamination of the normal tissue with the adjacent cancer). This type of observation is the same as what we see for the single cell WGBS data on mESC and MII cells; there is substantial heterogenity in global methylation for mESC and not for MII and we are concerned that our method fails on MII cells as evident by the low percentage of variance explained by the first eigenvector. More work is needed to firmly establish whether this ranking holds true for most primary tissues or might be a consequence of oncogenesis, manipulation in culture or a kind of unappreciated batch effect affected a well defined compartment of the genome. We note that the cause of the ranking does not matter; as long as the ranking is present it can be exploited to reconstruct A/B compartments.

The functional implications of A/B compartments have not been comprehensively described; we know they are associated with open and closed chromatin [Lieberman-Aiden et al., 2009], replication timing domains [Ryba et al., 2010, Pope et al., 2014], changes during mammalian development and are somewhat associated with gene expression changes [Dixon et al., 2015]. Our work makes it possible to more comprehensively study A/B compartments, especially in primary samples. We have illustrated this with a brief analysis of the relationship between A/B compartments and somatic mutation rate in prostate adenocarcinoma.

## Acknowledgments

Thanks to John Muschelli who made our Obs/Exp normalization function a thousand times faster. The results shown here are in whole or part based upon data generated by the TCGA Research Network: http://cancergenome.nih.gov/.

## Materials and Methods

### Infinium HumanMethylation450 BeadChip

We use the standard formula 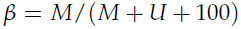 for estimating percent methylation given (un)methylation intensities 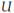 and *M*. Traditionally, the term M-value is used for the logit transform of the beta value, and we do the same.

With respect to CpG density, the 450k array probes fall into 4 categories that are related to CpG islands. CpG Island probes (30.9% of the array) are probes located in CpG islands, shore probes (23.1%) are probes within 2 kbs of CpG islands, and shelf probes (9.7%) are probes between 2 kbs and 4 kbs from CpG islands. Open sea probes (36.3%) are the rest of the probes. We use the term CpG resort probes to refer to the union of island, shore and shelf probes; in other words non-open sea probes.

### Methylation Data

Also see Table 3.

**Table 3.** Methylation data sources.

| Dataset | Cell type | n | Platform | Acc. | Reference |
| --- | --- | --- | --- | --- | --- |
| 450k-Fibroblast | Fibroblast (skin) | 62 | 450k | GSE52025 | Heyn et al. [2013] |
| 450k-EBV | LCL (EBV) | 288 | 450k | GSE36369 | Wagner et al. [2014] |
| 27k-EBV Vancouver | LCL (EBV) | 180 | 27k | GSE27146 | Fraser et al. [2012] |
| 27k-EBV London | LCL (EBV) | 160 | 27k | GSE26133 | Bell et al. [2011] |
| 450k-PRAD Normal | Prostate (normal) | 49 | 450k | TCGA | TCGA |
| 450k-PRAD Tumor | Prostate (tumor) | 340 | 450k | TCGA | TCGA |

**450k-Fibroblast dataset**: The study contains 62 samples from primary skin fibroblasts from Wagner et al. [2014]. The raw data (IDAT files) are available on GEO (Accession number: GSE52025)

**450k-EBV dataset**: The study contains 288 samples from EBV-transformed lymphoblastoids cell lines (LCL) [Heyn et al., 2013] from three HapMap populations: 96 African-American, 96 Han Chinese-American and 96 Caucasian. The data are available on GEO (Accession number: GSE36369).

**27k-EBV Vancouver**: The study contains 180 samples from EBV-transformed lymphoblastoid cell lines (LCL) [Fraser et al., 2012] from two HapMap populations: 90 individuals from Northern European ancestry (CEU), and 90 individuals from Yoruban (West African) ancestry (YRI). The processed data are available on GEO (Accession number: GSE27146)

**27k-EBV London**: The study contains 77 EBV-transformed lymphoblastoid cell lines (LCL) assayed in duplicates [Bell et al., 2011]. Individuals are from the Yoruba HapMap population, and 60 of them are also part of the 27k-EBV Vancouver dataset. The raw data (IDAT files) are available on GEO (Accession number: GSE26133)

**450k-PRAD Normal, 450k-PRAD Tumor**: At the time of download, the dataset contained 340 prostate adenocarcinoma tumor samples from The Cancer Genome Atlas (TCGA) [TCGA] along with 49 matched normal samples. We used the Level 1 data (IDAT files) available through the TCGA Data portal.

**PMDs-IMR90**: The partially methylated domain (PMD) boundaries from IMR90 [Lister et al., 2011] are available at http://neomorph.salk.edu/ips_methylomes/data.html.

**EBV hypomethylation blocks:** Hypomethylated blocks between EBV-transformed and quiescent B-cells were obtained form a previous study [Hansen et al., 2014]. Only blocks with a family-wise error rate equal to 0 were retained (see the reference).

### Processing of the methylation data

For the 450k-Fibroblast and 450k-PRAD datasets, we downloaded the IDAT files containing the raw intensities. We read the data into R using the illuminaio package [Smith et al., 2013]. For data normalization, we use the minfi package [Aryee et al., 2014] to apply the noob background subtraction and dye-bias correction [Triche et al., 2013] followed by functional normalization [Fortin et al., 2014b]. We have previously shown [Fortin et al., 2014b] that functional normalization is an adequate between-array normalization when global methylation differences are expected between individuals. For the 450k-EBV dataset, only the methylated and unmethylated intensities were available, and therefore we did not apply any normalization. For the 27k-EBV London dataset, IDAT files were available, and we applied the noob background correction and dye-bias correctin as implemented in the methylumi package [Triche et al., 2013]. For the 27k-EBV Vancouver dataset, IDAT files were not available and therefore we used the provided quantile normalized data as discussed in Fraser et al. [2012].

For quality control of the samples, we used the packages minfi and shinyMethyl [Aryee et al., 2014, Fortin et al., 2014a] to investigate the different control probes and potential batch effects. All arrays in all data sets passed the quality control. After normalization of the 450k array, we removed 17,302 loci that contain a SNP with an annotated minor allele frequency greater than or equal to 1% in the CpG site itself or in the single-base extension site. We used the UCSC Common SNPs table based on dbSNP 137. The table is included in the minfi package.

For the analysis of the 27k array data, we only considered probes that are also part of the 450k array platform (25,978 probes retained in total) and applied the same probe filtering as discussed above.

### Construction of 450k correlation matrices

For each chromosome, we start with a *p× n* methylation matrix *M* of *p* normalized and filtered loci and *n* samples. We use M-values as methylation measures. We compute the *p× p* matrix of pairwise probe correlations *C* = cor(*M*^*′*^), and further bin the correlation matrix *C* at a predefined resolution *k* by taking the median correlation for between CpGs contained in each of two bins. Because of the probe design of the 450k array, some of the bins along the chromosome do not contain any probes; these bins are removed. As discussed in the Results section, the correlations of the open sea probes are the most predictive probes for A/B compartments, and therefore the correlation matrix is computed using only those probes (36.3% of the probes on the 450k array). The inter-chromosomal correlations are computed similarly.

### Hi-C Data

Samples are described in Table 4.

**Table 4.**
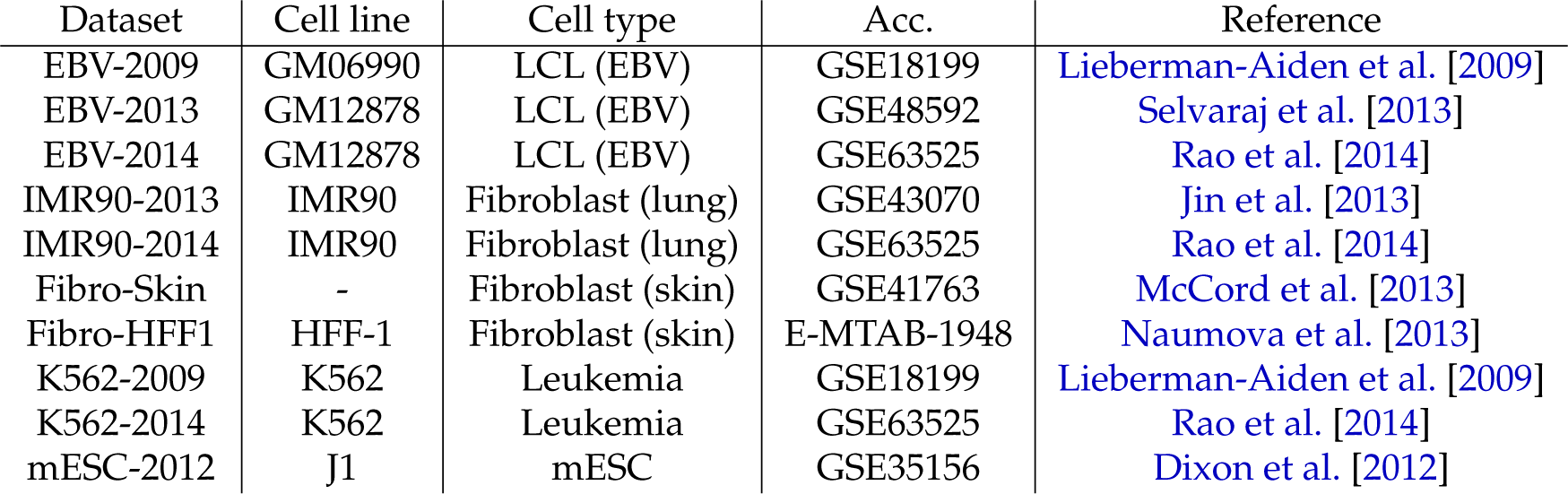
Hi-C data sources.

### Processing of the Hi-C data

For the Hi-C datasets EBV-2014, K562-2014 and IMR90-2014 from Rao et al. [2014], we used the raw observed contact matrices that were constructed from all read pairs that map to the human genome hg19 with a MAPQ ≥ 30. These contact matrices are available in the supplementary files of the GEO deposition (GSE63525). For the IMR90-2013 dataset from Jin et al. [2013], we used the online deposited non-redundant read pairs that were mapped with Bowtie [Langmead et al., 2009] to human genome hg18 using only the first 36 bases. For the EBV-2009 and K562-2009 datasets from Lieberman-Aiden et al. [2009], we used the mapped reads deposited on GEO (GSE18199). Reads were mapped to human genome hg18 using Maq, as described in [Lieberman-Aiden et al., 2009]. For the Fibro-Skin dataset from McCord et al. [2013], we merged the reads from two individuals with normal cells (Father and Age-Matched control). We used the processed reads of the GEO deposition (GSE41763) that were mapped using Bowtie2 to the hg18 genome in an iterative procedure called ICE previously described in Imakaev et al. [2012]. For the mESC-2012 dataset, we used the mapped reads deposited on GEO (GSE35156); reads were mapped to the mm9 genome.

For the EBV-2012 dataset from Selvaraj et al. [2013] and the Fibro-HFF1 dataset from Naumova et al. [2013], we downloaded the SRA experiments containing the FASTQ files of the raw reads. We mapped each end of the paired reads separately using Bowtie to the hg18 genome with the –best mode enabled. We kept only paired reads with both ends mapping to the genome.

For all datasets but the Hi-C datasets from Rao et al. [2014], we used the liftOver tool from UCSC to lift the reads to the human genome hg19 version for consistency with the 450k array. Reads from Rao et al. [2014] were already mapped to the hg19 genome.

### Construction of Hi-C matrices

As a first step, we build for each chromosome an observed contact matrix *C* at resolution *k* whose (*i*, *j*)’th entry contains the number of paired-end reads with one end mapping to the *i*-th bin and the other end mapping to the *j*-th bin. The size of the bins depends on the chosen resolution *k*. We remove genomic bins with low coverage, defined as bins with a total count of reads less than 10% of the total number of reads in the matrix divided by the number of genomic bins. This filtering also ensures that low mappability regions are removed.

To correct for coverage and unknown sources of biases, we implemented the iterative correction procedure called ICE [Imakaev et al., 2012] in R. This procedure forces bins to have the same experimental visibility. We apply the normalization procedure on a chromosome basis and noted that for each Hi-C dataset, the iterative normalization converged in less than 50 iterations. For the purpose of estimating A/B compartments, we further normalize the genome contact matrix by the observed-expected procedure Lieberman-Aiden et al. [2009], where each band of the matrix is divided by the mean of the band. This procedure accounts for spatial decay of the contact matrix.

### DNase-Seq data

Also see Table 5.

**Table 5.** DNase-Seq data sources.

| Dataset | Cell line | n | Acc. | Reference |
| --- | --- | --- | --- | --- |
| DNase-EBV | LCL (EBV) | 70 | GSE31388 | Degner et al. [2012] |
| DNase-IMR90 | Fibroblast (lung) | 4 | GSE18927 | Bernstein et al. [2010] |

**Table 6.**
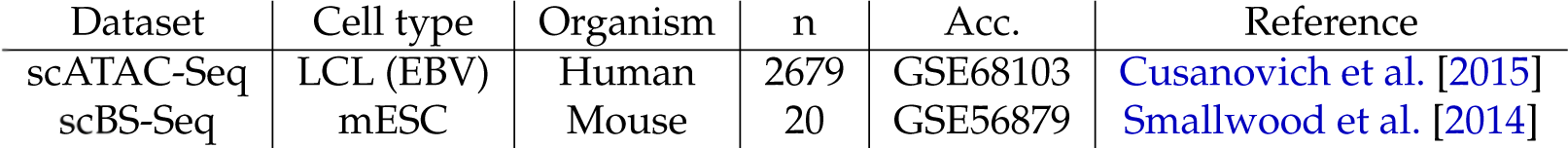
Single-cell epigenetic data sources.

**DNase-EBV dataset**: The study contains 70 biological replicates of EBV-transformed lymphoblastoid cell lines (LCL) [Degner et al., 2012] from the HapMap Yoruba population. The data are deposited on GEO (GSE31388) and raw files are available through http://eqtl.uchicago.edu/dsQTL_data/RAW_DATA_HDF5/.

**DNase-IMR90 dataset**: The dataset is composed of 4 technical replicates of the IMR90 fetal lung fibroblast cell line available on GEO (GSE18927).

### Processing of the DNase-Seq data

For the DNase-EBV dataset from Degner et al. [2012], we downloaded the raw reads in the HDf5 format for both the forward and reverse strands. We converted the reads to bedGraph, lifted the reads to the hg19 genome and converted the files to bigWig files using the UCSC tools. For the DNase-IMR90 dataset, we used the raw data already provided in the bigWig format. Reads were mapped to the hg19 genome. For both datasets, data were read into R by using the rtracklayer package [Lawrence et al., 2009]. We normalized each sample by dividing the DNase score by the total number of reads to adjust for library size.

### Construction of DNase signal and correlation matrices

For each sample, we construct a normalized DNase signal at resolution 100kb by taking the integral of the coverage vector in each bin. This was done using BigWig files and the rtracklayer package in R [Lawrence et al., 2009]. All DNAse datasets have the same read length within experiment (EBV/IMR90). This results in a *p* × *n* signal data matrix where *p* is the number of bins for the chromosome, and *n* the number of samples. The average DNase signal is the across samples mean of this matrix. The DNase correlation matrix is the *p* × *p* Pearson correlation matrix of the signal matrix.

### Single-cell ATAC-seq data

Single-cell ATAC-seq data was obtained from NCBI GEO using accession number GSE68103 described in Cusanovich et al. [2015]. We used data processed by the authors, specifically the file `GSM1647124 CtlSet1.dhsmatrix.txt.gz`. This experiments represents data on a mixture of two cell lines: Gm12878 and HL60. We use data processed by the authors of the paper, which consists of a matrix of accessibility across 195882 known hypersensitive sites (from ENCODE) and 4538 cells. Each hypersensitive site is furthermore characterized as being specific to GM12878, specific to HL60 or common across the two cell types. To classify each cell to a cell type, we computed the total number of reads in each of the cell type specific hypersensitive sites, normalized by the total number of reads in all hypersensitive sites for that cell (arbitrarily scaled by 2000). These numbers are displayed in Figure 20. We visually selected the line displayed in this figure (*y* = *x* − 3) to separate out cells from the GM12878 cell type. Subsequently we discarded hypersensitive sites which had no reads in any of the cells and obtained 632 bins at a resolution of 100kb on chromosome 14. Eigenvectors were computed and smoothed as described below.

### Single-cell WGBS data

Single-cell whole genome bisulfite sequencing data was obtained from NCBI GEO using accession number GSE56879 described in Smallwood et al. [2014]. We used data processed by the authors, specifically the files `GSM1370555 Ser X.CpG.txt.gz` where `X` takes value 1 to 20. These files describe the single CpG methylation levels of 20 individual cells for mouse embryonic stem cells cultured in serum conditions. We removed CpGs within 4kb of a CpG Island (using the CpG Islands defined in Wu et al. [2010]), as we did for the 450k methylation array data. We next binned the genome in 100kb bins and computed, for each bin, the average methylation value across all CpGs in the bin. Bins with a total coverage of less than 100 was removed from analysis. This resulted in a binned methylation matrix which was used to compute an empirical correlation matrix. Eigenvectors were computed and smoothed as described below.

### Eigenvector analysis

To obtain eigenvectors of the different matrices from Hi-C, DNA methylation and DNase data, we use the non-linear iterative partial least squares (NIPALS) algorithm implemented in the mixOmics package in R [Dejean et al., 2014]. Each eigenvector is smoothed by a moving average with a 3-bin window, with the following exceptions. For the 450k data, we used two iterations of the moving average smoother. For the single cell epigenetic data we used a window size of 5 bins with two iterations of the moving average smoother for ATAC-seq and 3 iterations for WGBS.

When we compare eigenvectors from two different types of data, we only consider bins which exists in both data types; some bins are filtered out in a data-type dependent manner for example because of absence of probes or low coverage. This operation slightly reduces the number of bins we consider in each comparison.

Because the sign of the eigenvector is arbitrarily defined, we use the following procedure to define a consistent sign across different chromosomes, datasets and data types. For Hi-C data and DNase data, we correlate the resulting eigenvector with the eigenvector from Lieberman-Aiden et al. [2009]; changing sign if necessary to ensure a positive correlation. For DNA methylation data, we use that the long-range correlations are significantly higher for the closed-closed interactions. We therefore ensure that the eigenvector has a positive correlation with the column sums of the binned correlation matrix; changing sign if necessary. This procedure results in positive values of the eigenvector being associated with closed chromatin and the ‘B’ compartment as defined in Lieberman-Aiden et al. [2009] (in this paper they ensure that negative values are associated with the closed compartment).

To measure the similarity between two eigenvectors, we use two measures: correlation and compartment agreement. The correlation measure is the Pearson correlation between the smoothed eigenvectors. The compartment agreement is defined as the percentage of bins that have the same eigenvector sign, interpreted as the percentage of bins that belong to the same genome compartment (A or B) as predicted by the two eigenvectors. Occasionally, this agreement is restricted to bins with an absolute eigenvector value greater than 0.01 to discard uncertain bins.

Because open chromatin regions have very high DNase signal in comparison to closed chromatin regions, the DNase signal distribution is highly skewed to the right; therefore we center both the average signal and the first eigenvector by subtracting their respective median, before computing the correlation and agreement.

### Somatic mutations in PRAD

We obtained a list of somatic mutations in PRAD from the TCGA data portal (https://tcga-data.nci.nih.gov/tcga/). Several lists exists; we used the Broad Institute curated list (`broad.mit.edu_IlluminaGA_curated_DNA_sequencing level2.maf`. To obtain capture regions, we queried the CGHub website (https://cghub.ucsc.edu) and found that all samples were profiled using the same capture design described in the file `whole_exome_agilent_1.1_refseq_plus_3_boosters.targetIntervals.bed` obtained from the CGHub bitbucket account.

Somatic mutation rates in each 100kb genomic bin were computed as the number of mutations inside each bin, divided by the length of the capture regions inside the bin.

### Software

Methods for performing the analysis of 450k methylation arrays described in this manuscript will be added to the minfi package [Aryee et al., 2014].

